# Low fidelity DNA polymerase IV accelerates genome evolution in *Pseudomonas aeruginosa*

**DOI:** 10.1101/2024.10.22.619742

**Authors:** Sofía D. Castell, Consuelo M. Fernandez, Ignacio N. Tumas, Lucía M. Margara, María C. Miserendino, Danilo G. Ceschin, Roberto J. Pezza, Mariela R. Monti

**Author notes:** Joint Authors (Consuelo M. Fernandez, Ignacio N. Tumas).

## Abstract

Specialized DNA polymerases are crucial for bypassing lesions and facilitating various cellular processes. Despite extensive research, the mutagenic effects of these error-prone enzymes on genomes are still not fully understood. In this study, we examined the genomic instability caused by Pol IV in the bacterial pathogen *Pseudomonas aeruginosa*. Pol IV mutagenesis primarily involved the misincorporation of oxidized guanine nucleotides opposite template adenine. This activity led to a distinctive mutational signature, characterized by A to C transversions occurring preferentially at AT sites flanked by a 5’G and/or 3’C. Furthermore, Pol IV preferentially targeted specific chromosomal locations near the replication termination region and rRNA-encoding operons. Several genes associated with virulence, motility, antibiotic resistance and chemotaxis are located in these difficult-to-replicate regions, making them hotspots for Pol IV-mutagenesis. Notably, half of the mutation events catalyzed by Pol IV effectively impaired gene function. This can be attributed to the strong bias of Pol IV for mutating specific codons with its preferred sequence contexts, leading to substitutions primarily to the unreactive Ala and Gly residues. Remarkably, mutation signatures identified for Pol IV were also present in genomes from clinical isolates of *P. aeruginosa*, providing compelling evidence for its role in genetic diversification during pathogen adaptation.

## INTRODUCTION

Mutations are prominent sources of genetic diversity, significantly impacting adaptation, evolution, senescence, and tumorigenesis. While often viewed as a stochastic process, there is increasing evidence that mutations can also exhibit strong biases^1,2^. These biases have been linked to mutagenic mechanisms at play and various genomic features that influence mutation processes, such as DNA sequence contexts, transcription levels, and replication timing^1–3^. Different mutational processes produce distinctive combinations of mutation types and specific context-dependent patterns, referred to as mutational signatures. Specific mutational signatures have been associated with environmental mutagen exposures and DNA repair defects^1–3^. Thus, identifying the mutagenic processes underlying a mutational signature is essential for predicting adaptive outcomes in pathogenic bacteria and guiding therapeutic strategies for genetic diseases.

Errors made by DNA polymerases (Pol) during replication are significant contributors to mutagenesis and genomic instability. High-fidelity replicative Pols carry out most of the DNA synthesis required for genome duplication^4^. These enzymes achieve accurate replication by discriminating between correct nucleotides during polymerization and correcting errors through proofreading^4,5^. In addition, overall fidelity is maintained by the post-replicative mismatch repair (MMR)^6^. Mismatches uncorrected by proofreading are recognized by the MutS repair protein, which subsequently recruits MutL to remove the newly synthesized strand containing the incorrect nucleotide.

Faithful DNA replication by replicative Pols can be challenged upon encountering template lesions or difficult-to-replicate regions, such as those with secondary structures or physical impediments^7^. Since stalled replication forks can threaten cell viability, replicative Pols rely on specialized translesion synthesis (TLS) Pols to continue DNA replication past damaged bases or challenging genomic areas^4,5,7^. TLS Pols inherently exhibit lower accuracy compared to replicative Pols due to their lack of intrinsic proofreading activity and a more open active site architecture^4,5^. Consequently, their ability to relieve replication stall often comes at the price of increased mutagenesis.

Most organisms possess a wide repertoire of TLS Pols dedicated to specific roles^4,5,7^. Among these, Pol IV is the only DNA polymerase conserved across all life domains^8^. The mutagenic potential of this Y-family DNA Pol has been well-established *in vitro*. Both Pol IV and its eukaryotic homolog, Pol κ, facilitate the erroneous bypass over oxidized DNA templates and the incorrect incorporation of oxidized nucleotides^9–13^. These TLS Pols also catalyze low-fidelity replication of undamaged DNA and efficiently extend misaligned primer/template termini^14,15^. Additionally, Pol IV plays a significant role in promoting genetic diversity under stressful conditions in bacteria. Its error-prone synthesis is particularly relevant for mutagenesis during stationary-phase and under the SOS DNA-damage response^16–19^. This activity is also linked to the development of antibiotic resistance and virulence in pathogenic bacteria^20–22^. In human cancer, mutational loads have been associated with the activity of Pol κ, which appears to generate mutation clusters reminiscent of the stress-induced mutagenesis response observed in bacteria^23^. While Pol IV is crucial for stress-induced mutagenesis, it does not significantly contribute to genetic diversity under normal growth conditions. The loss of Pol IV does not affect the rate or spectra of genomic mutations in unstressed bacterial cells^16,24–29^, suggesting a tight regulation of Pol IV-promoted mutagenesis under normal conditions that breaks down during stress.

Several regulatory mechanisms limit the DNA synthesis by low-fidelity TLS Pols^4,30^. A key regulatory point involves their interaction with processivity sliding clamps, the prokaryotic β clamp and the archaeal/eukaryotic proliferating cell nuclear antigen (PCNA)^4,30^. These ring-shaped proteins provide a topological link to the DNA template and localize TLS Pols to DNA replication sites^31^. The association with clamps, facilitated by clamp binding motifs in Pols and hydrophobic clefts in the clamps, is essential for effective functioning of TLS Pols^26,32^. In this context, our previous research revealed a non-canonical function of MutS in modulating the interaction of Pol IV with β clamp^26^. The repair protein can displace Pol IV from the clamp by competing for binding to the hydrophobic cleft on the processivity factor. According to our model, Pol IV can effectively alleviate stalling of the replication fork; however, if it incorporates an erroneous nucleotide, MutS recognizes the mispair and disrupts the Pol IV-β clamp interaction. This action by MutS restricts further replication by Pol IV and promotes the removal of the incorrect nucleotide through the proofreading activity of a high-fidelity Pol. Similar mechanisms for coordinating the access of TLS Pols to DNA replication sites by MutS has also been observed in eukaryotes. In human cells, the MutS homolog, MutSα, regulates Pol κ- and Pol η-dependent TLS by promoting monoubiquitination of PCNA in response to UV and oxidative damage, respectively^33,34^. This transient post-translational modification of PCNA facilitates recruitment of the relatively error-free TLS polymerase.

Although Pol IV is a key player in adaptive mutagenesis, its influence on mutagenesis across the entire genome remains poorly understood. To address this knowledge gap, we examined the Pol IV-mutational profile at the genomic level and its impact on evolutionary processes in the human opportunistic pathogen, *Pseudomonas aeruginosa*. We focused our study on this organism because adaptive processes in cystic fibrosis chronic pulmonary infections represent one of the best-studied models of long-term microbial evolution within a human host. *P. aeruginosa* gradually transitions from an acute virulent pathogen in early stages to a host-adapted pathogen in chronic lung infections^35,36^. This adaptive process is primarily mediated by inactivating mutations that turn off acute virulence factors, such as motility appendages and pigments, while enhancing traits that promote long-term survival^35–37^. These evolutionary changes are believed to be driven in response to selective pressures, including the highly oxidative environment present the cystic fibrosis lung ecosystem^35,37^.

## MATERIALS AND METHODS

### Bacterial strains and culture media

The experiments were carried out using *Pseudomonas aeruginosa* PAO1 and *mutT*::ISphoA/hah (location phoAwp01q3F06) MPAO1 strains. The *mutT* mutator strain was obtained from the University of Washington Genome Center^38^. Transposon insertion within the corresponding gene was confirmed by PCR analysis following the manufacturer’s instructions. For the PAO1 and *mutT* MPAO1 strains, isogenic *mutS*^β^, *dinB* and *mutS*^β^ *dinB* derivatives were made by allelic exchange using the mobilizable suicide plasmid pKNG101 as previously described^26^. The *mutS*^β^ strain harbors a chromosomal *mutS* allele, named *mutS*^β^, which codifies a MutS mutant unable to interact with β clamp as the putative β clamp binding ^816^QSDLF^820^ motif was changed to ASDAA. The *dinB* strain harbors a chromosomal *dinB* allele containing a 753 bp deletion at the 5’ of the 1050 bp open reading frame. Luria-Bertani or M9 minimal media were used. To prepare inocula, bacteria were cultured on agar plates from frozen stocks and sub-cultured in liquid LB medium overnight with shaking at 200 rpm at 37°C.

### Estimation of mutation rates by fluctuation assays

Mutation levels of different target genes on the *P. aeruginosa* chromosome were measured by estimating mutation rates to resistance to ciprofloxacin (*nfxB* is mutated at the ciprofloxacin concentration used in this work), rifampicin (*rpoB* is mutated) and amikacin (several target genes are associated to amikacin resistance). Mutation rates were determined by the modified Luria-Delbruck fluctuation test^39^. Independent cultures were obtained as follows: cells were inoculated in LB liquid medium (∼500 cells/ml) and grown to exponential phase. For complementation assays, cells freshly transformed with the derivatives of p5BAD were inoculated in LB medium containing 5 μg/ml gentamicin. For paraquat (methyl viologen)-induced mutagenesis, cells were cultured in LB medium amended with 100 µM paraquat. Cell death was negligible under these conditions, and thus, fluctuation assays were conducted. After cultures reached late-exponential phase (∼0.1-1.0 × 10^9^ cells/ml), aliquots from successive dilutions were plated onto LB agar or LB agar containing 5 μg/ml gentamicin to determine the number of viable cells, and onto LB containing 0.5 or 1.3 μg/ml ciprofloxacin, 50 μg/ml rifampicin or 2.5 or 10 μg/ml amikacin to select resistant cells. Colonies were scored after 24-48 hours. The rSalvador program^40^ was used to calculate maximum likelihood estimates of m under the Lea-Coulson model^41^, and μ and 95% confidence limits; and to perform the statistical comparison of fluctuation assay data based on the likelihood ratio test.

To estimate mutation rates in cells growing on LB solid media, freshly streaked colonies were grown on LB plates. The number of viable and resistant cells per colony was determined by suspending colonies, diluting and plating onto LB and LB amended with antibiotics, respectively. Each colony was treated as an individual culture in a fluctuation test and the mutation rate was calculated as previously described.

As a control, the minimal inhibitory concentration (MIC) for the antibiotics used in the fluctuation assays was determined according to the guidelines of the Clinical and Laboratory Standards Institute. All strains showed comparable MICs, confirming that the changes in mutation rates were not owing to differences in the MIC.

### *nfxB* inactivation

Inactivation of *nfxB* was analyzed using a luminescent reporter for detection of the transcriptional de-repression of the *mexCD-oprJ* operon^42^. The transcriptional fusion of the *mexC* promoter to the *luxCDABE* reporter operon was inserted into the attTn7 site of the *P. aeruginosa* chromosome using the pUC18-mini-Tn7T-Gm-lux vector^43^. Chromosomal insertions were confirmed by PCR as recommended in the published protocol.

Ciprofloxacin resistant clones were obtained from exponential cultures grown as described in fluctuation assays. Independent clones (25-30) selected at ciprofloxacin 0.5 μg/ml were picked up onto LB agar plates and then onto LB agar containing ciprofloxacin to confirm that exhibited a stable phenotype. Luminescence was measured in exponentially growing cells as follows: cultures in LB liquid medium were diluted to an OD 600 nm of 0.1 and seeded in 96-well plates (100 μl). The 96-well plate was then imaged by integrating the luminescence signal for 20 s in a NightOWL LB 983 instrument, and the number of photons emitted per well was quantified using the Berthold WinLight 32 Software. The fold of increase in the luminescence intensity for the ciprofloxacin resistant clones was calculated as the ratio of the number of photons exhibited by cultures of the ciprofloxacin resistant clones and the parental strain.

### Mutation accumulation (MA) experiments

MA lines were initiated from single colonies isolated from *mutT*, *mutT mutS*^β^, *mutT dinB* and *mutT mutS*^β^ *dinB* strains. The founder colonies were obtained by streaking from a freezer stock onto LB agar plates and incubating the plate overnight at 37°C. Eight independent well-isolated colonies from each founder strain were streaked out on LB plates and incubated at 37 °C for 24 h. This single-cell bottleneck was repeated 100 times. The colony chosen for passage was the one closest to the end of the streak trace to ensure random selection of colonies. After each transfer, the original plate was stored as a backup plate at 4 °C. If a well-isolated colony was not available for streaking on a particular day, a new streak was made from another colony on the stored plate. The MA experiment was repeated twice. 8-14 MA lines of each founder strain were subsequently used for whole genome sequencing.

### Genomic DNA sequencing and variant calling

For whole-genome sequencing of MA lines, a single colony from the last passage of each MA was inoculated in liquid LB and grown overnight with shaking. This culture was used to perform a freezer stock and to extract genomic DNA using the GenElute Bacterial Genomic DNA purification kit (Sigma-Aldrich). Genomic DNA was also harvested from each ancestor strain for sequencing in order to analyze the single-nucleotide polymorphisms present prior to initiating the MA experiment protocol and filter them from the analysis. Sequence data were obtained from 150-bp, paired-end reads using the Illumina NovaSeq 6000 (Novogene, USA). The mean ± standard deviation of the average depth of coverage across all samples was 190 ± 14.

Raw sequencing data underwent preprocessing using Fastp (version 0.23.2)^44^. This step involved cleaning adaptors, removing bad sequences, and filtering out low-quality sequences to ensure the reliability of subsequent analyses. Cleaned fastq files were aligned to the reference genome using SubRead (version 2.0.1)^45^. Variant Calling was performed using the Genome Analysis Toolkit (GATK) pipeline (version 4.2.5)^46,47^. The BAM file containing aligned reads underwent several steps, including sorting, modifying the read groups, marking duplicates, reordering, and indexing^48^. Annotation of the identified variants was performed using snpEff (version 5.1)^49^. The *P. aeruginosa* PAO1 genome (NCBI Reference Sequence, NC_002516.2; https://www.pseudomonas.com/strain/show?id=107) served as the reference for mapping the variants and providing biological information for interpretation and analysis. This annotation process involved mapping the variants to specific genes, functional regions, and associated biological information, facilitating their interpretation and analysis.

The genome sequence of the founder *mutT* strain differs from PAO1 at 26 single-nucleotide polymorphisms (Table S11). *mutT mutS*^β^, *mutT dinB* and *mutT mutS*^β^ *dinB* have additional mutations to that found in the *mutT* strain (Table S11). The identity of each MA line was confirmed by examining the presence of the *mutS*^β^ mutation and the *dinB* internal deletion in the *mutS* and *dinB* genes, respectively. For quality control, we verified that mutations specific to founder strains were also called correctly in all descendant MA lines. A few MA lines shared some identical mutations, suggesting cross-contamination during passaging. One of the pair of suspect strains was randomly chosen to be discarded from the analysis to ensure each replicate line in the dataset was unique. The mutation dataset was fitted to Poisson distributions. The generalized Pearson statistic and model deviance were used to assess Poisson goodness of fit.

For analysis of *P. aeruginosa* whole-genomes from clinical and environmental isolates, complete genomes were obtained from www.pseudomonas.com/strain and aligned to the reference PAO1 genome using MiniMap2 (version 2.28)^50^. Variant calling was conducted using Bcftools (version 1.15.1)^51^. Annotation of the identified variants was performed using snpEff (version 5.1)^49^.

### Estimation of generations and mutation rates from MA experiments

For each endpoint MA line and the ancestors, the total number of viable cells per colony was determined by excising colonies from agar plates, re-suspending in LB and plating dilutions on LB agar plates. The number of generations per growth cycle was estimated as the log2 of total cell numbers. The number of viable cells per colony was similar between MA lines and the corresponding ancestor. Only one MA line derived from the *mutT mutS*^β^ *dinB* strain showed a decrease in the number of viable cells. This MA line was discarded from the analysis. The mean number of generations per MA line was 2655 (SD= 51), 2639 (SD= 109), 2631 (SD= 58) and 2673 (SD= 64) for *mutT*, *mutT mutS*^β^, *mutT dinB* and *mutT mutS*^β^ *dinB*, respectively.

Mutation rates per nucleotide site per cell division were calculated by dividing the total number of accumulated mutations by the total number of generations that the MA lines passed and by the appropriate number of sites (A:T sites, G:C sites, etc.). Confidence limits (CLs) for the mutation rates were calculated from the mean and variance of the mutations per MA line for each strain, using the critical values of the t (student) distribution^28^. The two-way ANOVA test was performed to compare mutation rates between strains after determining normality via the Shapiro-Wilks test and homogeneity of variance via the Levene test.

### Phenotypic characterization of MA lines

Overnight cultures in liquid LB were started from individual colonies of each strain in triplicate. For swimming motility, 2 µl of a 1/10 dilution of overnight cultures was inoculated into the center of Tryptone swim plates (1% tryptone, 0.5% NaCl, 0.3% agar). Plates were incubated at 37°C for 24 hours. Motility was assessed by scanning the plates and measuring the diameter of the circular turbid zone using ImageJ. For pyoverdine quantification, overnight cultures were adjusted to 10^6^ cells per ml in M9 minimal medium and incubated with shaking at 200 rpm at 37°C for 16 hours. A 1.5 mL culture was centrifuged at 13,000 rpm for 2 min. 200 μL cell-free supernatants were transferred to a clear-bottom black-side 96-well plate and fluorescence (excitation 360/40 nm; emission 460/40 nm) was measured using a BioTek Synergy HT equipment. Pyoverdine levels were expressed as fluorescence normalized by the biomass of cultures, measured by absorbance (600 nm).

## RESULTS

### Pol IV mutagenesis is associated with the misincorporation of oxidized nucleotides

We previously established that *P. aeruginosa* MutS interaction with β clamp is critical for controllig Pol IV mutagenic replication in exponentially growing cells^26^. We showed that Pol IV contributes to spontaneous mutagenesis in a *mutS*^β^ PAO1 mutant strain, in which the chromosomal *mutS* allele was replaced by *mutS*^β^ that encodes a MutS version unable to bind to β clamp^52^, but not in the parental (WT) PAO1 strain. Briefly, the *mutS*^β^ strain exhibited a high proportion of AT>CG base substitutions in the reporter chromosomal *nfxB* gene, which encodes a negative regulator of the *mexCD-oprJ* genes for drug efflux^53^. Deletion of the Pol IV encoding *dinB* gen (*mutS*^β^ *dinB* strain) significantly decreased the rate of this transversion. Conversely, inactivation of *dinB* did not change the mutation *nfxB* spectra in the WT background (*dinB* strain).

AT>CG transversions can result from the incorporation of 8-oxo-2’-deoxyguanosine-5’-triphosphate (oxodGTP) opposite a template adenine, a process Pol IV has been shown to catalyze *in vitro*^12,13,54^. Therefore, we hypothesized that MutS might regulate Pol IV mutagenesis associated with the erroneous insertion of oxidized nucleotides. To test this, we first analyzed the mutagenesis under oxidative damage by estimating mutation rates in cells treated with the free-radical producing agent paraquat (Figure 1A and S1A, Table S1 and S2). We found that *mutS*^β^ cells exposed to a non-lethal paraquat concentration showed a 2-fold increment in mutation rates to resistance to ciprofloxacin (Cip^r^) and amikacin (Amk^r^) relative to untreated cells. In contrast, treatment with paraquat did not affect the mutation rates in the *mutS*^β^ *dinB*, WT and *dinB* strains.

**Figure 1.**
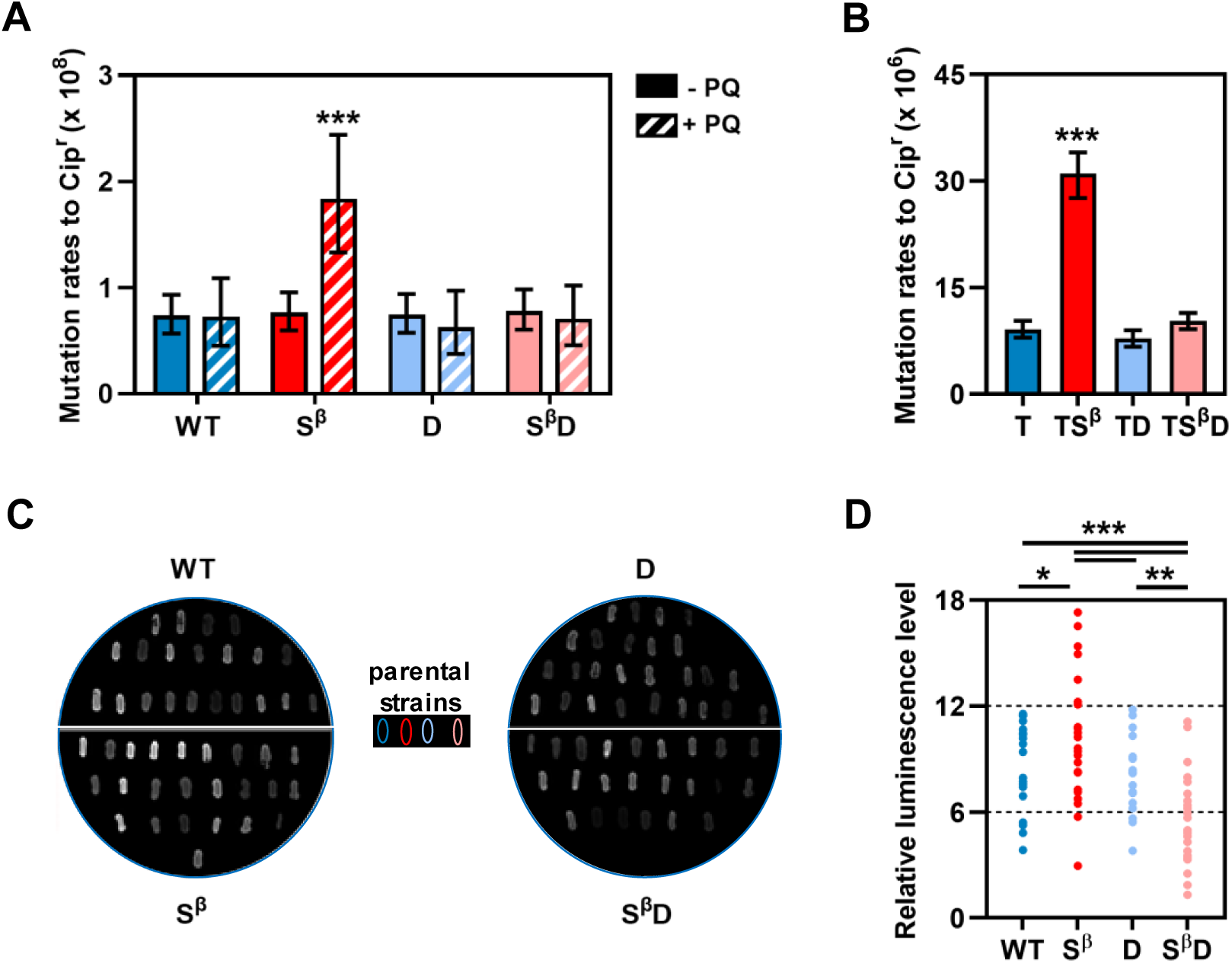
Contribution of Pol IV to oxodGTP mutagenesis and gene inactivation. (A) Mutation rates for the WT, *mutS*^β^ (S^β^), *dinB* (D) and *mutS*^β^ *dinB* (S^β^D) strains treated (+ PQ) or not (-PQ) with paraquat. (B) Mutation rates for the *mutT* (T), *mutT mutS*^β^ (TS^β^), *mutT dinB* (TD) and *mutT mutS*^β^ *dinB* (TS^β^D) strains. Mutation rates to ciprofloxacin resistance (Cip^r^) per replication and 95% confidence limits were calculated as described in Materials and Methods. Error bars represent the upper and lower 95% confidence limits. Data were statistically analyzed using the likelihood ratio test. (C) Representative images of luminescent Cip^r^ clones from the WT, S^β^, D and S^β^D strains carrying the chromosomal transcriptional fusion of the *mexCD*-*oprJ* promoter to the *luxCDABE* operon. (D) NfxB inactivation in Cip^r^ clones from the WT, S^β^, D and S^β^D strains. Luminescence intensity was measured in exponential cultures of the individual Cip^r^ clones. The relative luminescence level was calculated as the ratio between the luminescence intensity in the Cip^r^ clone and the corresponding parental strain. Values are the mean from three independent experiments. Results were evaluated using one-way ANOVA and Tukey tests.

We next estimated spontaneous mutation rates in *mutT* mutant cells (Figure 1B and S1B, Table S3). The MutT enzyme hydrolyzes oxodGTP to prevent its incorporation into DNA^55^. Thus, *mutT* deficiency should increase oxodGTP cellular levels and promote its DNA incorporation by Pol IV. We estimated mutation rates in exponentially growing *mutT*, *mutT mutS*^β^, *mutT dinB* and *mutT mutS*^β^ *dinB* cells. Consistent with previous data^56^, the *mutT* strain showed a strong mutator phenotype, exhibiting two to three orders of magnitude higher mutation rates relative to WT. *mutT mutS*^β^ displayed a 3- to 5-fold increased mutation rates to resistance to rifampicin (Rif^r^), Cip^r^ and Amk^r^ relative to *mutT*. The Pol IV-deficient strain, *mutT mutS*^β^ *dinB*, exhibited a significant decrease in mutation rates compared to *mutT mutS*^β^, reaching values comparable to those observed in the *mutT* strain. No significant differences in mutation rates were detected among the *mutT* and *mutT dinB* strains.

To confirm that Pol IV is involved in the increased mutagenesis in *mutT mutS*^β^, we performed complementation assays by introducing arabinose-inducible p5BAD plasmids bearing the wild-type and mutant *dinB* alleles into the *mutT mutS*^β^ *dinB* strain (Figure S1C, Table S4). Pol IV expression from plasmid p5BAD-*dinB* increased mutation rates 2-fold relative to control cells harboring the empty vector p5BAD. Mutation rates in cells carrying the p5BAD-*dinB*D8A derivative, encoding the inactive DNA polymerase Pol IV-D8A mutant^25^, failed to enhance mutation rates.

In summary, our findings demonstrate that Pol IV-mediated mutagenesis involves the erroneous incorporation of oxidized nucleotides into DNA, and this activity is regulated by MutS through its interaction with β clamp.

### Pol IV generates highly inactive variants of the reporter *nfxB* gene

Loss-of-function mutations are a common adaptive strategy in *P. aeruginosa* during cystic fibrosis airway infections^35,37^. Specifically, inactivating mutations in coding sequences involved in antibiotic resistance such as *nfxB* are frequently selected^57^. Taking this into account we ascertained whether Pol IV could produce such loss-of-function mutations by evaluating the repressor activity of NfxB variants obtained from the WT, *mutS*^β^, *dinB* and *mutS*^β^ *dinB* strains (Figure 1C, 1D and S1D). Inactivating mutations within *nfxB* impair or abolish the NfxB repressor activity, leading to overexpression of the MexCD-OprJ efflux pump and resistance to ciprofloxacin^53,58^. To measure the repressor activity of NfxB variants, we integrated a luminescence-based reporter construct, containing the transcriptional fusion of the *mexC* promoter to the *luxCDABE* operon, in the attTn7 chromosomal site of the PAO1 strains^42^. The *luxCDABE* operon is normally repressed at a basal level by endogenous NfxB, however, loss-of-function mutations in *nfxB* de-repress *luxCDABE* expression^58^.

Independent Cip^r^ clones were obtained from exponential cultures of WT, *mutS*^β^, *dinB* and *mutS*^β^ *dinB* strains. The degree of NfxB inactivation in these Cip^r^ clones was quantified as the fold-increase in the luminescence intensity relative to the corresponding parental strain (Figure 1C, 1D and S1D). Clones showing luminescence increases of up to 6-, 12- and 18-fold were arbitrarily categorized as expressing NfxB variants with low, medium and high inactivation levels, respectively. Cip^r^ clones from *mutS*^β^ expressed NfxB variants showing a high (29%) or medium (63%) inactivation level. These high inactivated NfxB variants were not detected in *mutS*^β^ *dinB*, which showed Cip^r^ clones with a medium (39%) and low (61%) inactivation level. In the WT strain, 81% of Cip^r^ clones exhibited medium luminescence levels while 19% showed low levels. A similar proportion was observed in the *dinB* strain, indicating no significant difference between them. These results suggest that Pol IV efficiently generates highly inactive NfxB variants in the *mutS*^β^ genetic background. As expected from the fact that Pol IV does not contribute to spontaneous mutagenesis in the WT strain, there were no differences in the kind of variants detected in the WT and *dinB* strains.

### Impact of Pol IV activity on spontaneous mutations throughout the genome

#### The AT>CG transversion dominates the mutation spectra in the mutT strains

To assess the mutator activity of Pol IV at the genome-wide level, we conducted mutation accumulation experiments. This involved propagating eight independent lines derived from *mutT*, *mutT mutS*^β^, *mutT dinB* and *mutT mutS*^β^ *dinB* founder strains through repeated single-individual bottlenecks achieved by streaking for single colonies (Figure 2A). This mutation accumulation experiment was repeated twice. Under these conditions, mutations should accumulate in a neutral manner with negligible selective pressure^59^. After approximately 2,650 generations, we identified accumulated mutations by sequencing the genome of 8-14 lines from both replicates and the corresponding ancestor to an average coverage of 190X. We anticipated observing Pol IV mutagenic activity in the *mutT mutS*^β^ background as MutS is unable to regulate Pol IV binding to the processivity β clamp factor. Such mutagenesis should not be observed in the *mutT* background, where the MutS control mechanism is operating. Conversely, predicting the effect of Pol IV deficiency in *mutT dinB* and *mutT mutS*^β^ *dinB* is challenging, as the mutagenic outcome depends on which alternative mutagenic processes prevail in the absence of this DNA polymerase.

**Figure 2.**
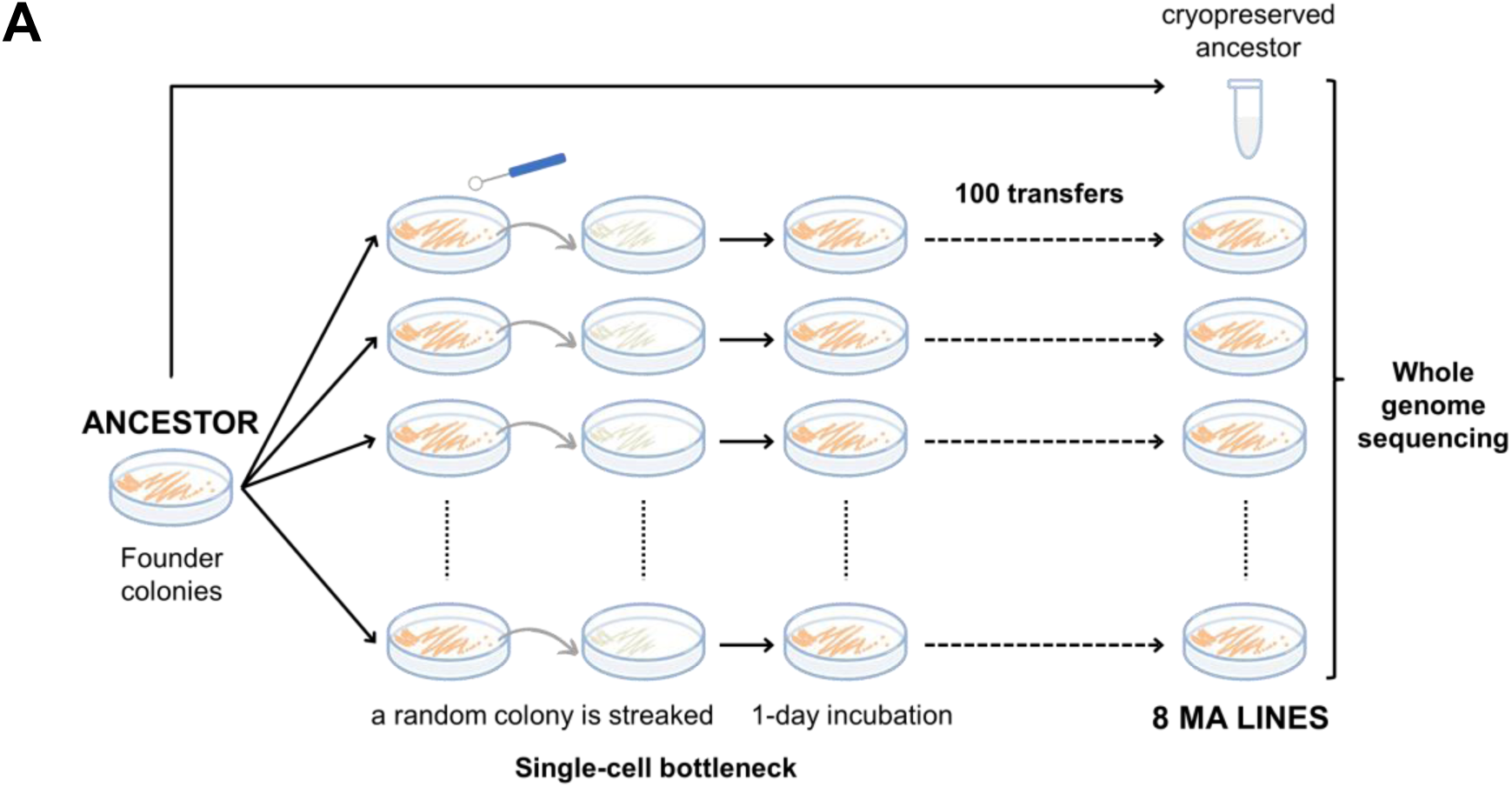
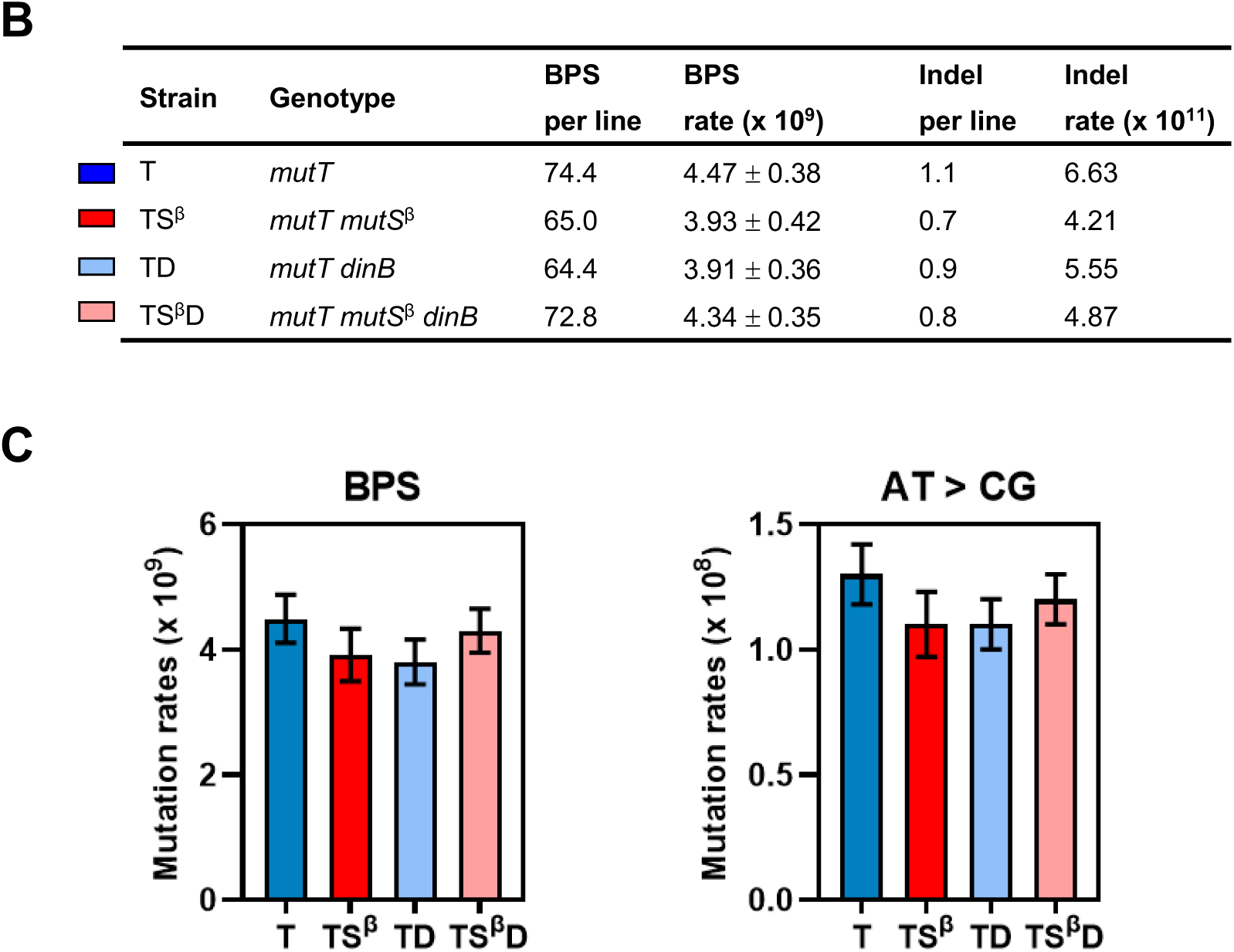
Spontaneous mutagenesis in the genome of the *mutT*, *mutT mutS*^β^, *mutT dinB* and *mutT mutS*^β^ *dinB* strains. (A) Experimental strategy of the mutation accumulation (MA) experiment. (B) Strain name, genotype, and mutation rates from MA experiments. (C) Total BPS and AT>CG mutation rates. Mutation rates per generation and nucleotide (genome: 6,264,404; AT base pairs: 2,092,311) and 95 % confidence limits were calculated as described in Materials and Methods. Confidence limits for values with fewer than 20 mutation events were not calculated due to the lack of statistical power. Plotted data and error bars represent the mutation rate and 95% confidence limits. The data were statistically analyzed using the two-way ANOVA and Tukey tests.

Before the initiation of the mutation accumulation experiments, and as the physiological state of the cells changes when growing in colonies compared to liquid cultures from fluctuation assays, we estimated mutation rates in colonies to analyze Pol IV mutagenesis under these conditions (Figure S2 and Table S5). Compared to mutagenic values of the *mutT* strain, the *mutT mutS*^β^ strain exhibited a significant 12- and 7-fold increase in the rates to Cip^r^ and Amk^r^, respectively. There were no significant differences in mutation rates to Rif^r^ between the *mutT* and *mutT mutS*^β^ strains. Deletion of *dinB* decreased the mutation rate to Cip^r^ in *mutT mutS*^β^, indicating that Pol IV is involved in the *nfxB* mutagenesis, but had no effect on the rate to Amk^r^ and Rif^r^. These data suggest that Pol IV mutagenesis is locus specific in the *mutT mutS*^β^ strain growing in colonies. Conversely, spontaneous mutations in the *mutT* strain appear independent of Pol IV, as its deficiency did not affect mutation rates.

The mean number of mutations per line was 74.4, 65.0, 64.4 and 72.8 for the *mutT*, *mutT mutS*^β^, *mutT dinB* and *mutT mutS*^β^ *dinB* strains, respectively (Figure 2B). Analysis of the pooled mutation datasets revealed a fit to Poisson distributions (χ^2^ = 6.2, *df* = 3, *P* = 0.10), indicating consistent mutation rates throughout the mutation accumulation experiments. The spectrum of all strains was dominated by base-pair substitutions (BPSs) (Figure 2B and 2C). The BPS mutation rate did not differ significantly between the four groups with different founder genotypes (p = 0.42), averaging 4.16 × 10^−9^ BPSs per nucleotide per generation. This value is 53-fold higher than the genome-wide BPS mutation rate in the PA14 WT strain^60^ and represents a mild increase compared to the 266-fold higher mutation rate of the PA14 *mutS* strain^60^. The BPS spectrum across all four strains was strongly biased toward the AT>CG transversion (Figure 2C and Table S6), accounting for approximately 97% of the BPSs with an average rate of 1.22 × 10^−8^ (p = 0.33, indicating no significant differences among the four strains). This aligns with expectations based on previous locus-specific experiments in *P. aeruginosa*^58^ and genome-wide analysis in *Escherichia coli*^27^. Nearly all the lines accumulated a single small insertion/deletion (indel, 1-79 bp), occurring at an average rate of 5.32 × 10^−11^ indels per nucleotide per generation (Figure 2B). This rate value is close to that observed in the PA14 WT strain^60^. In conclusion, a faulty hydrolysis of oxodGTP exclusively increases the AT>CG substitution rate (561-fold, *mutT* PAO1 vs PA14 WT strains), impacting the AT sites in the *P. aeruginosa* genome. Remarkably, this mutational profile is conserved in the four *mutT* strains.

We next assessed selection efficiency during line propagation by examining the ratio of mutations in coding versus noncoding DNA (Table S7). The expected ratio of mutations in AT nucleotides between coding and noncoding regions in the PAO1 genome is 7.95 under neutral accumulation conditions. In the *mutT*, *mutT mutS*^β^ and *mutT mutS*^β^ *dinB* strains, the observed ratio was not statistically different from this expectation. The *mutT dinB* strain showed a lower ratio due to an increased proportion of mutations in noncoding regions. This fact is inconsistent with a strong role for selection as selective pressure is assumed to be minimal for noncoding regions. Selective pressure was further evaluated by the ratio of the number of nonsynonymous to synonymous substitutions (Table S7). Under the assumption that synonymous mutations are relatively neutral, and given the codon usage in PAO1^61^, the expected ratio is 17.47. For all strains, the ratio of nonsynonymous to synonymous mutations did not significantly deviate from this random expectation. These findings collectively support that mutations accumulate in a nearly neutral fashion during the mutation accumulation protocol.

#### Pol IV is involved in the mutagenesis of specific functional pathways

We asked whether specific functional pathways are preferentially affected by Pol IV-mutagenic activity. With this aim, we first classified all pools of randomly sampled mutations in coding regions (approximately 430-670 per founder strain) into the 27 functional categories of the PseudoCAP database^62^. The mutation rate per AT nucleotides in coding regions per generation was estimated for each PseudoCAP category in the four strains (Table S8). These rates were compared with the whole-genome mutation rate in coding regions for each strain to identify deviations (Figure 3A). We observed a higher relative mutation rate among genes categorized under “antibiotic resistance and susceptibility”, “chemotaxis” and “motility and attachment” in the *mutT mutS*^β^ strain. Conversely, an equal or lower rate was detected for these categories relative to the whole-genome rate in the *mutT*, *mutT dinB* and *mutT mutS*^β^ *dinB* strains. Within these functional categories, we found an enrichment of the following pathways in *mutT mutS*^β^: Che and Chp chemosensory pathways; structural components of the type IVa pili; structural components of the flagellum and transcriptional regulators of flagellar genes; and the ATP binding cassette (ABC) and resistance/nodulation/cell division (RND) efflux pumps. Moreover, the genes encoding the outer membrane porins and cytotoxins within the “membrane proteins” and “secreted factors” categories, respectively, were highly represented in *mutT mutS*^β^ but not in *mutT*, *mutT dinB* and *mutT mutS*^β^ *dinB*. Mutation rates in these pathways were 2- to 5-fold higher in *mutT mutS*^β^ compared to that observed in *mutT*, *mutT dinB* and *mutT mutS*^β^ *dinB*, with exception of the flagellar genes that also showed a high mutation rate in *mutT mutS*^β^ *dinB* (Figure 3B). On average, 73% of *mutT mutS*^β^ lines accumulated mutations in these pathways (Figure S3). Notably, *mutT mutS*^β^ lines showed up to 4 mutations in a single pathway, averaging 6 mutations per line across these six pathways. In contrast, 29% of lines from the other strains accumulated mutations with an average of 2.3 mutations per line. In addition to these six pathways, the pyoverdine biosynthesis and secretion genes were also a target of Pol IV-mutagenesis. Mutation levels in this pathway were similar for all strains (Figure 3C and S3), however, subsequent data in this study indicated that Pol IV is responsible for mutations found in these genes in the *mutT mutS*^β^ lines. Furthermore, 41 genes associated with other pathways were consistently mutated in the *mutT mutS*^β^ lines, but not in *mutT*, *mutT dinB* and *mutT mutS*^β^ *dinB*. Based on these findings, we identified 96 target genes of Pol IV mutation, comprising a total of 156 mutation events (Table S9). These Pol IV-target genes and mutations have been pivotal in characterizing the features of Pol IV-induced mutagenesis described in the following sections of this article.

**Figure 3.**
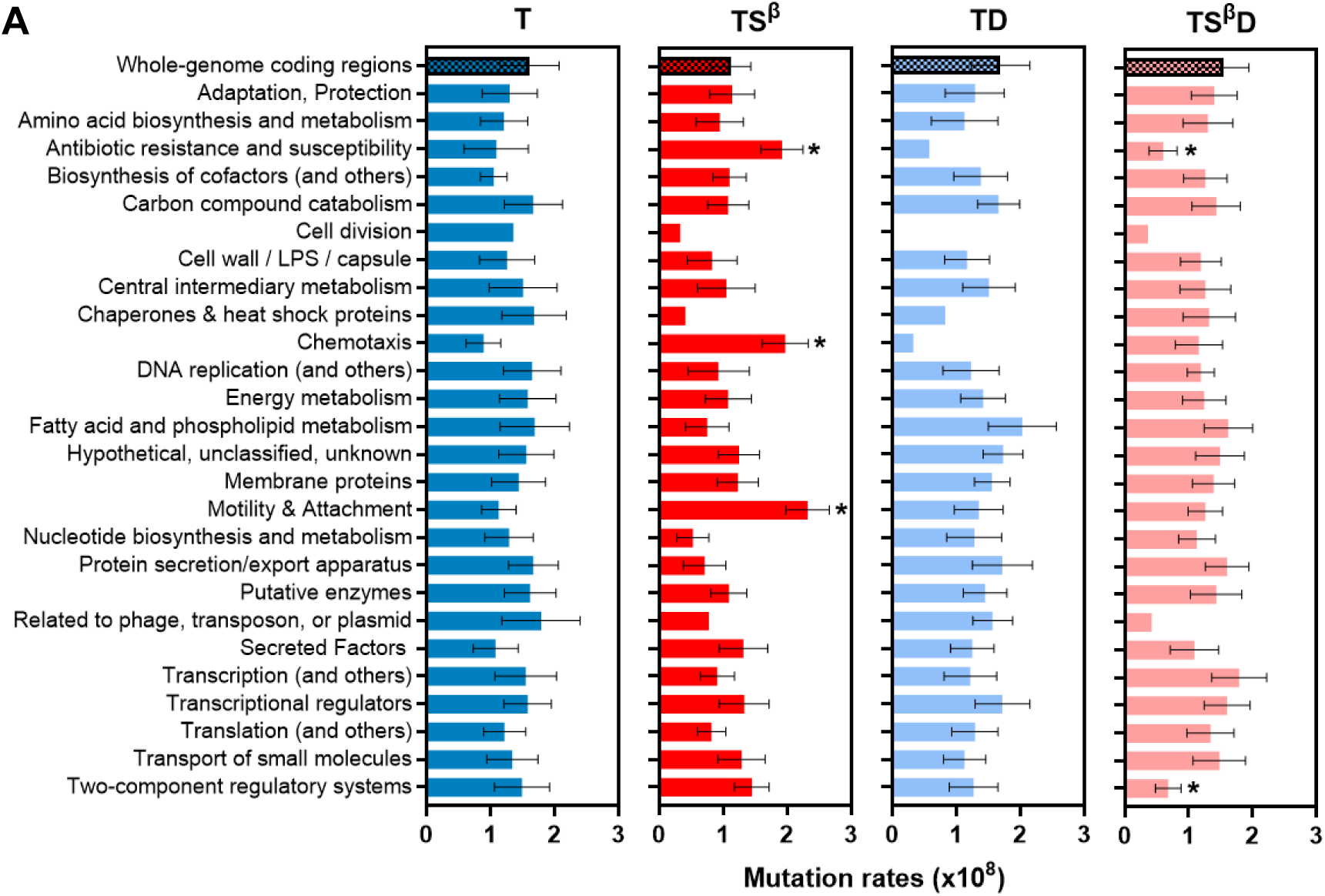
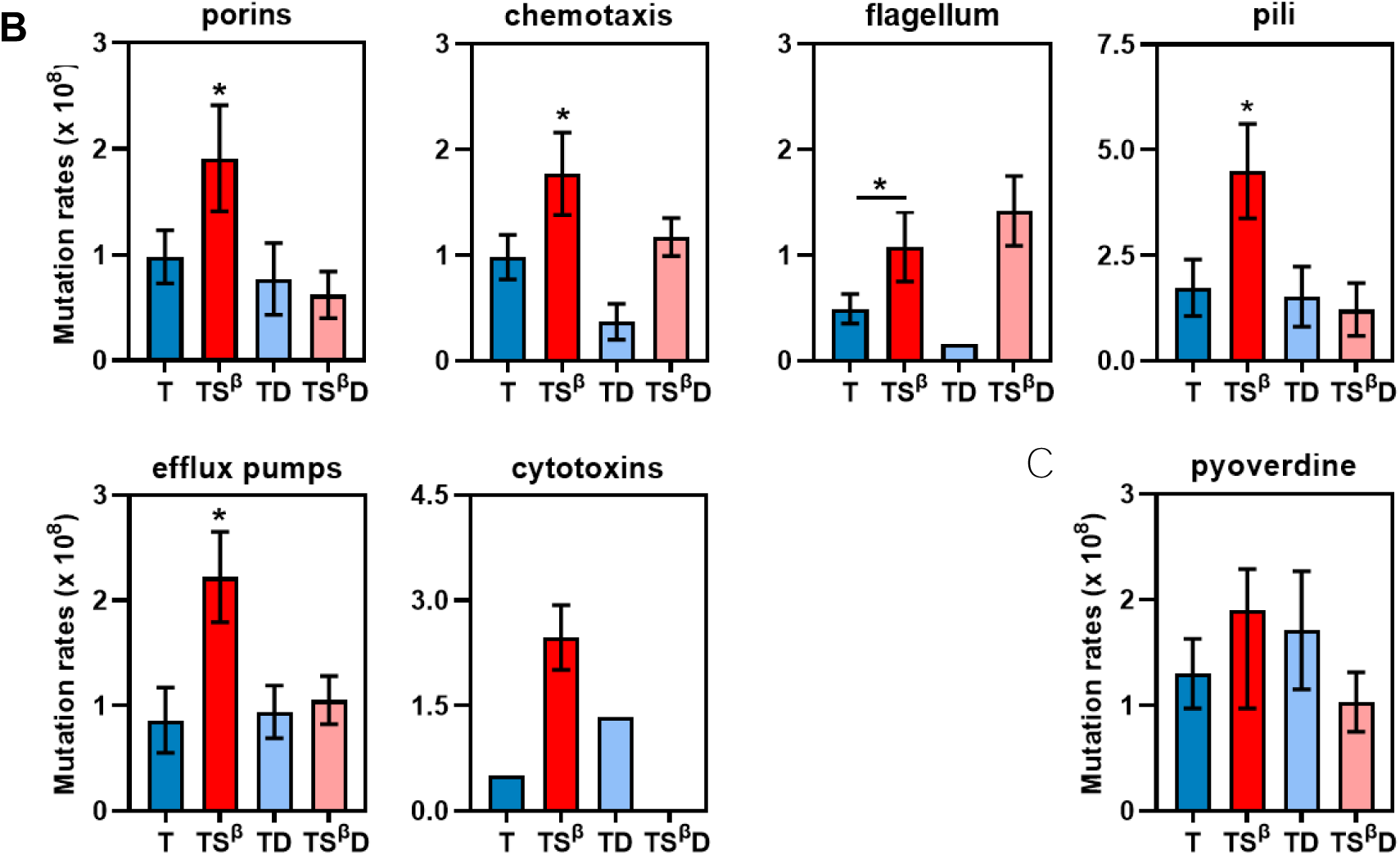
Pol IV displays a preference for mutating specific functional pathways. BPS mutation rates in PseudoCAP categories (A) and functional pathways (B) for the *mutT* (T), *mutT mutS*^β^ (TS^β^), *mutT dinB* (TD) and *mutT mutS*^β^ *dinB* (TS^β^D) strains. AT>CG mutation rates per generation and nucleotide, and 95% confidence limits were calculated as described in Materials and Methods. The number of AT in the coding regions of the genes included in each PseudoCAP category and functional pathway was used to estimate mutation rates. Confidence limits for values fewer than 5 BPS events were not calculated. Error bars represent the upper and lower 95% confidence limits. No overlap of 95% confidence intervals indicates statistically significant differences. In panel A, the asterisks show significant differences with the mutation rates in the whole-genome coding regions for each strain.

#### Pol IV mutagenesis is biased by the local sequence context

Previous studies using reporter genes have shown that mutations induced by Pol IV are context dependent. Specifically, this DNA polymerase promotes deletions at mononucleotide repeats and base substitutions at the G**X**C trinucleotide (**X** represents the mutated base)^24,26,63–65^. Throughout this report, the mutated base is indicated in bold and DNA sequences are presented in the 5′-to-3′ direction. We searched DNA sequence preferences surrounding mutation target sites for Pol IV using various approaches. Initially, the 5’ and 3’ bases flanking the mutated nucleotide and their reverse complements were recorded for the AT>CG transversions accumulated in the four strains (approximately 422-655 mutations per founder strain) and induced by Pol IV (151 mutations). Context-dependent mutation rates for mutated AT nucleotides at each of the 16 possible trinucleotides were estimated as the number of mutations at each trinucleotide normalized to the number of this trinucleotide in the genome and generations (Figure 4A and S4). Pol IV significantly promoted mutations at G**A**C+G**T**C and A**A**C+G**T**T trinucleotides, with mutation rates 2- and 3-times higher compared to those observed in the *mutT*, *mutT dinB* and *mutT mutS*^β^ *dinB* strains. The mutation rate at G**A**A+T**T**C, associated with oxodGTP mutagenesis^27^, was highest among mutations induced by Pol IV and accumulated in the strains. In contrast, A**A**A+T**T**T, A**A**T+A**T**T and T**A**A+T**T**A trinucleotides were not hotspots for Pol IV-induced mutagenesis. Mutations at these trinucleotides were not detected among the Pol IV-mutated genes or exhibited a 2-fold lower value than that observed for the strains.

**Figure 4.**
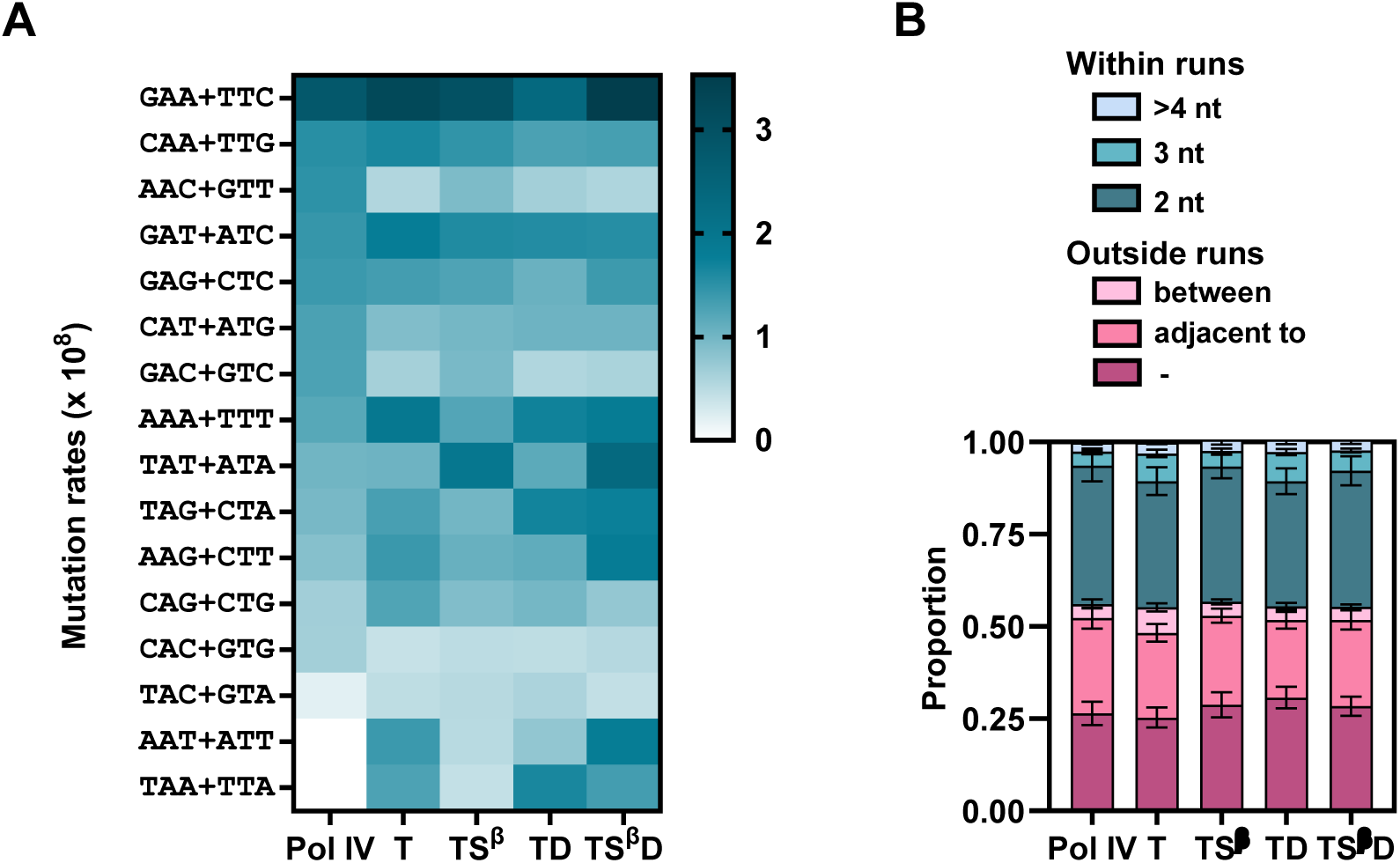
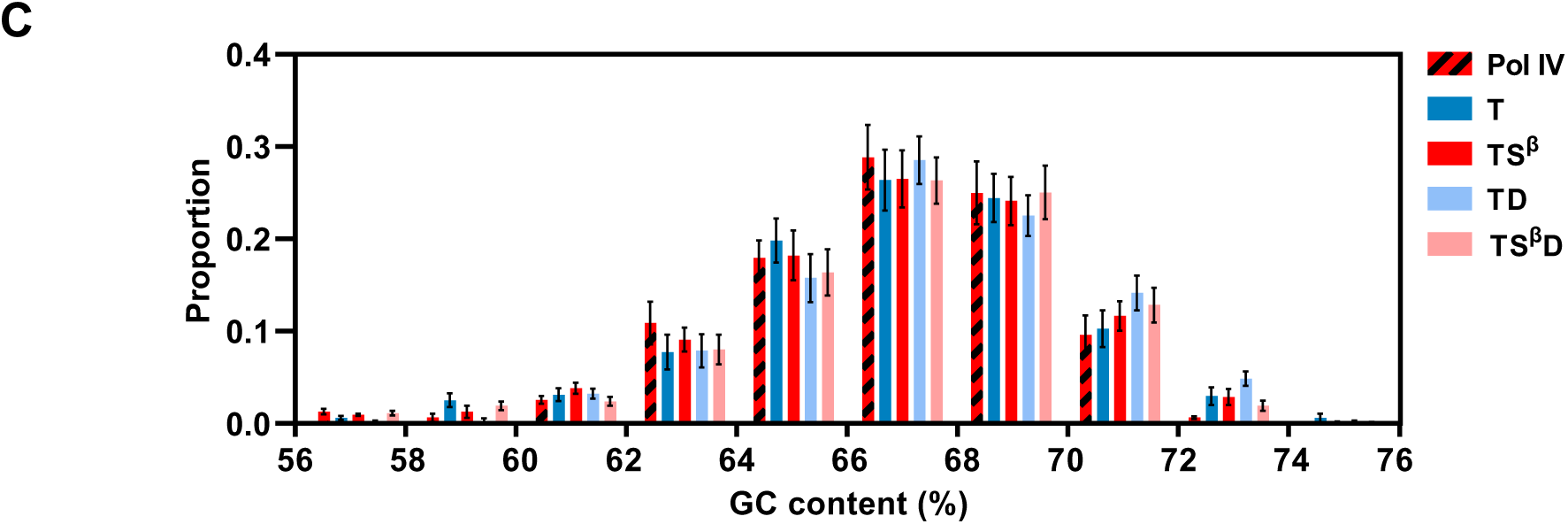
Pol IV-induced mutations are biased by the local sequence context. (A) BPS mutation rates at trinucleotides. Mutation rates per generation and trinucleotide were estimated for AT>CGs in the Pol IV-target genes and accumulated by the *mutT* (T), *mutT mutS*^β^ (TS^β^), *mutT dinB* (TD) and *mutT mutS*^β^ *dinB* (TS^β^D) strains. Triplets are written 5′ to 3′ with the target site in the center. (B) BPS at mononucleotide repeats. Data bar fill colors indicate the proportion of AT>CGs located outside, adjacent to, between two or within a mononucleotide repeat for mutations accumulated in the Pol IV-mutated genes and the four strains. (C) GC content of mutated genes. The proportion of AT>CG mutations was plotted against the GC content of the mutated genes. Error bars represent standard deviations. Statistical analysis was performed using the Kruskal–Wallis test.

We next explored the ±10 bp neighboring sequence to identify additional nucleotides influencing Pol IV-mutagenesis using the MEME motif detection algorithm^66^. This analysis did not reveal any conserved sequence context associated with the Pol IV-induced mutations.

We further investigated if mononucleotide repeats are hotspots for the AT>CG base substitutions promoted by Pol IV. By visual inspection of sequences surrounding mutated nucleotides, we categorized whether the mutated AT was located outside, adjacent to, between two, or within a mononucleotide repeat; and calculated the proportion of mutations at each context (Figure 4B). There were no significant differences in the location of mutations between Pol IV-induced mutations and mutations in the strains (p = 0.93). Approximately 44% of mutations occurred within mononucleotide repeats, mostly at runs of 2 nucleotides. The remainder of AT>CG mutations occurred adjacent to (23%) or were not linked to (28%) mononucleotide repeats.

Considering that mutation rates in genome sequences are dependent on local GC content^67,68^, we examined whether GC content influences Pol IV-mutagenesis in coding regions. To accomplish this, we analyzed the distribution of AT>CG transversions among mutated genes based on their GC content, which ranged from 56 to 76 % (Figure 4C). The distribution of AT>CG mutations in the Pol IV-target genes was indistinguishable from the overall datasets of the four strains (p = 0.97). Pol IV-target genes exhibited an average GC content of 66.6%, similar to that of mutated genes in the *mutT*, *mutT mutS^β^, mutT dinB* and *mutT mutS*^β^ *dinB* strains (66.7, 66.8, 67.3 and 66.7%, respectively), and consistent with coding regions in the PAO1 genome (66.6%). Our analysis of sequence features influencing Pol IV-mutagenesis revealed that this DNA polymerase prefers specific DNA sequence contexts. 5’ G and/or 3’ C surrounding the AT target base is a hotspot for Pol IV mutations at the genome level. This observation aligns with the fact that G**X**C is the preferred context observed in the reporter *nfxB* gene of *P. aeruginosa*^26^. Conversely, the presence of T and A bases adjacent to the mutation site or within mononucleotide repeats do not promote AT>CG mutations by Pol IV. Additionally, the GC content of the target sequence does not impact Pol IV-mutagenesis.

#### Pol IV mutations are concentrated at specific genome positions

Chromosomal locations could contain mutational hotspots for Pol IV that result in a non-random distribution of mutations along the length of the circular *P. aeruginosa* genome. To identify preferred genomic targets of Pol IV-mutagenesis, the PAO1 genome was divided into 25 bins of 0.25 Mbp and mutation rates were estimated for each bin (Figure 5). For mutation rate calculations, the number of AT>CG per bin was divided by the number of AT in coding regions within the bin and generations. The data of Pol IV-mutated genes and mutated genes across all strains are depicted separately for the right and left replichores in Figure 5B (see individual data in Figure S5A). A wave-like distribution of AT>CG mutations was observed for all strains, similar to patterns described for the PA14 WT and *mutS* strains^60^. However, there were no significant differences in mutation rates across the genome. Notably, Pol IV-induced mutations were not equally distributed along the genome. Three large clusters of mutations appeared at 2.50-2.75 Mbp near the replication terminus region (position ∼3.20 M)^69^ and at 5.00-5.25 and 5.50-5.75 Mbp close to the rRNA-encoding genes. In these regions, mutation rates were 4- and 3-fold higher, respectively, compared to those observed for the strains. A fourth cluster was noticed at 0.25-0.50 Mbp, showing a significant 2-fold increased mutation rate in the Pol IV-target regions compared to the strains. Interestingly, the majority of type IV pili genes (94%) and a significant proportion of genes involved in pyoverdine biosynthesis, chemosensory pathways, ABC/RND efflux pumps, and porins (20-70%) were located within these four chromosomal regions (Figure S5B). Interesting, Pol IV-induced mutations were sparsely detected in the genome regions spanning 3.00-3.75 Mbp and 5.75-6.25 Mbp.

**Figure 5.**
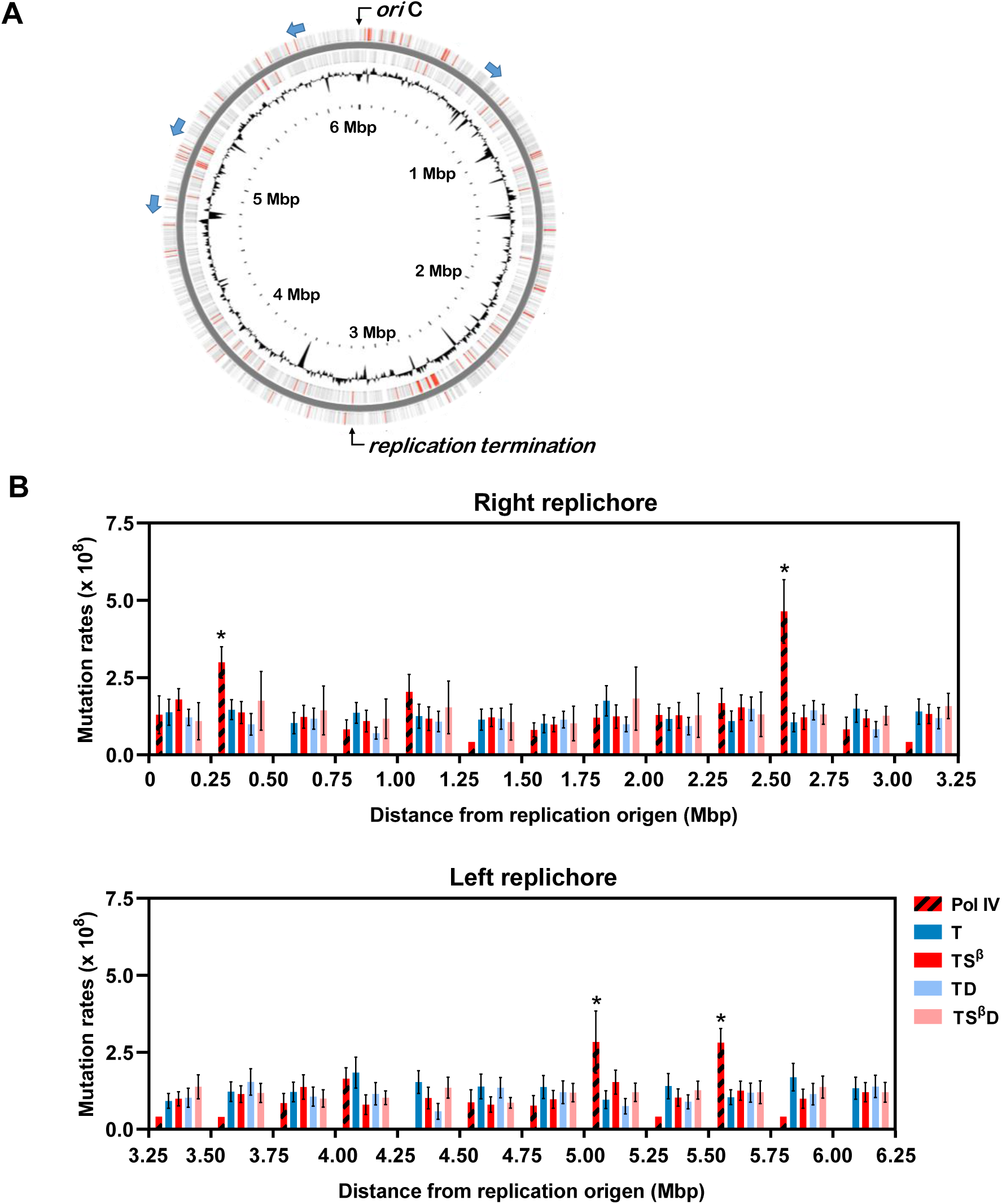
Pol IV-mutational hotspots in the *P. aeruginosa* genome. (A) Schematic of the PAO1 chromosome. The circular map displays the localization of the origin (*ori*C) and replication termination regions, ribosomal RNA clusters (blue arrows), genes (grey lines) and Pol IV-target genes (red lines). The outer and inner bands represent the plus and minus strands, respectively. The black plot shows the percentage GC content plotted as the average for non-overlapping 0.5-kb windows spanning one strand for the whole genome. The genomic map was generated by Proksee ^70^. (B) Mutation rates along the *P. aeruginosa* chromosome. AT>CG mutation rates were plotted as a function of distance from the origin of replication in 0.25-Mbp bins for the Pol IV-target genes and the *mutT* (T), *mutT mutS*^β^ (TS^β^), *mutT dinB* (TD) and *mutT mutS*^β^ *dinB* (TS^β^D) strains. Mutation rates were estimated as the number of AT>CGs per the number of AT in coding regions in each bin and generations. Error bars show 95% confidence limits. Confidence limits for values fewer than 5 BPS events were not calculated. No overlap of 95% confidence intervals indicates statistically significant differences.

These data indicate a biased distribution of Pol IV-induced mutations towards specific regions of the *P. aeruginosa* chromosome, with the 2.50-2.75 Mbp region exhibiting the highest Pol IV-mutagenic activity.

#### Highly inactive variants are generated by Pol IV in the mutation accumulation assay

We evaluated the ability of Pol IV to generate inactivating mutations under the non-selective conditions of our mutation accumulation experiment. To conduct this analysis, we characterized *mutT*, *mutT mutS*^β^, *mutT dinB* and *mutT mutS*^β^ *dinB* lines carrying mutations in genes involved in pyoverdine biosynthesis/secretion and flagellum synthesis, for their associated phenotypes. We analyzed these pathways because we had enough mutant lines to compare *mutT mutS*^β^ with the other strains. In lines with mutations in the pyoverdine pathway, we quantified the production and secretion of the siderophore in culture supernatants using fluorescence (Figure 6A). Among the 6 *mutT mutS*^β^ lines, 3 showed significantly decreased pyoverdine production, showing 20- to 40-fold lower pyoverdine levels compared to the *mutT mutS*^β^ ancestor. However, none of the 16 lines from *mutT*, *mutT dinB* and *mutT mutS*^β^ *dinB* exhibited reduced pyoverdin production, despite 6 of these lines had 2 mutations in the pyoverdine pathway. Thus, the 22 mutation events accumulated in these strains did not effectively inactivate the pyoverdine genes. Interestingly, 3 lines from *mutT* and *mutT dinB* displayed a hyperproduction phenotype, showing 8- to 13-times higher siderophore levels than the ancestors. For lines with mutations in flagella genes, we examined swimming motility (Figure 6B). Among the evolved lines from *mutT mutS*^β^, 50% effectively inactivated flagellum synthesis genes. Of 6 *mutT mutS*^β^ lines, 1 showed a complete absence of swimming motility and 2 exhibited decreased motility compared to the ancestor. Conversely, only 1 line (12%) showed reduced swimming motility, while no significant change was observed in the remaining 7 lines from *mutT*, *mutT dinB* and *mutT mutS*^β^ *dinB*. Among these lines, 4 had mutations in 2 to 3 flagella genes, accumulating a total of 12 mutations in the flagella pathway, none of which affected swimming motility. In summary, our data demonstrated that Pol IV is highly proficient at inactivating genes.

**Figure 6.**
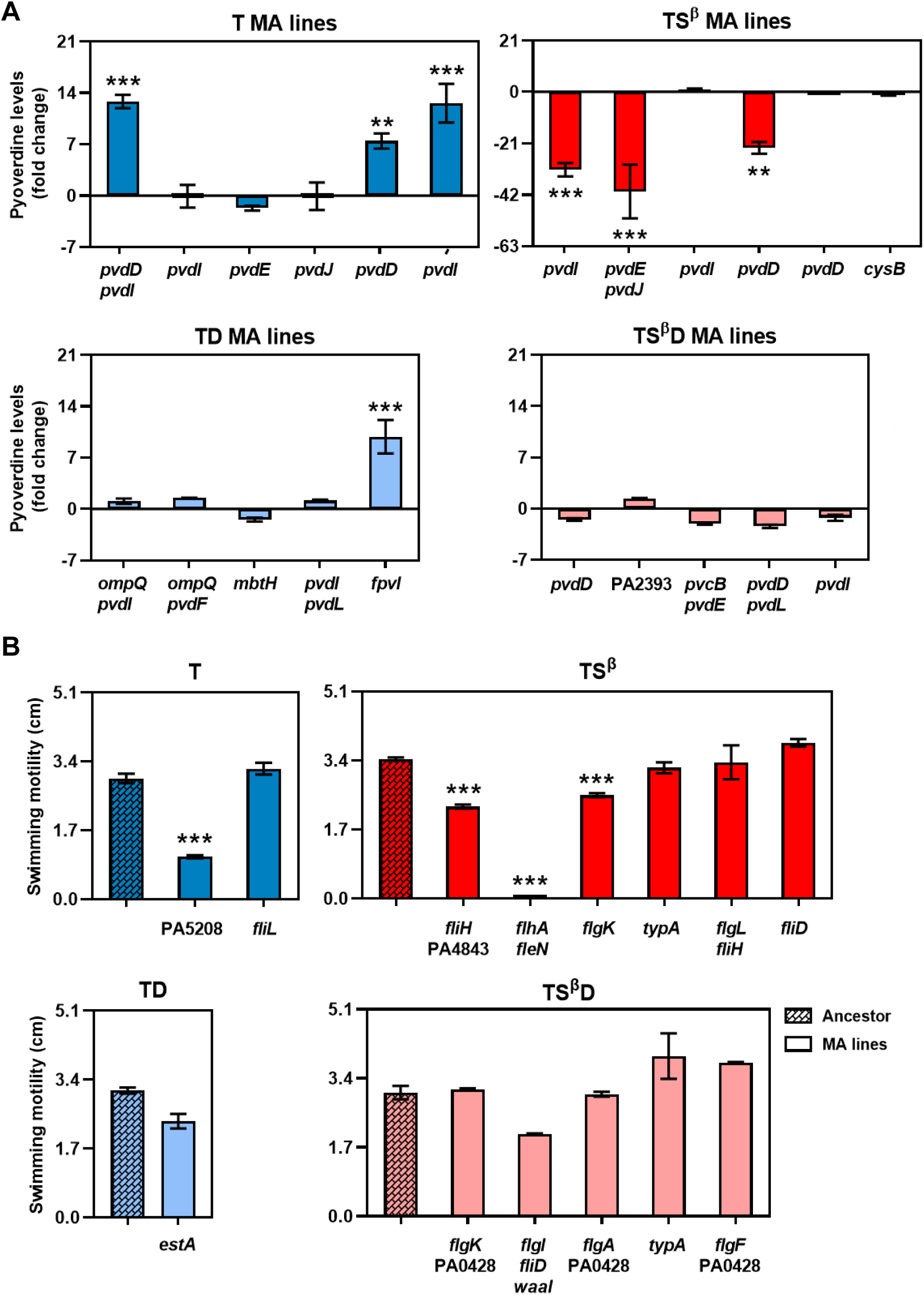
Pol IV generates highly inactive variants. (A) Siderophore production/secretion. Pyoverdine levels were quantified by fluorescence and normalized to bacterial growth. Bars show the pyoverdine fluorescence in cultures of the mutation accumulation (MA) lines relative to that observed for the corresponding ancestral strain. Error bars show standard deviations. (B) Swimming motility. Diameters across the zones of bacterial swimming were measured, and the means and standard deviation are shown. Data represent the average of three independent experiments. One-way ANOVA followed by Tukey tests were performed to compare each *mutT* (T), *mutT mutS*^β^ (TS^β^), *mutT dinB* (TD) and *mutT mutS*^β^ *dinB* (TS^β^D) line with the corresponding ancestral strain. Mutated genes from the pyoverdine and flagella pathways are indicated at the bottom of each MA line bar.

#### Pol IV preferentially mutates specific codons

We investigated the features of Pol IV-mutagenesis that led to effective gene inactivation. First, we analyzed whether Pol IV has a bias for mutating specific codons by estimating codon mutation rates. Mutation rates were calculated for each codon as the ratio between the number of a mutated codon and the number of that codon in the PAO1 genome and generations (Figure 7A and S6A). We noticed a strong preference of Pol IV for mutating specific codons. GAC, GTC, TCC, GTG and GAA codons showed ∼4-, 4-, 3-, 3- and 2-fold increased mutation levels in Pol IV-mutated genes than that observed for the *mutT*, *mutT dinB* and *mutT mutS*^β^ *dinB* strains. In agreement with this, a high proportion of Ser, Val, Asp and Glu were mutated among the Pol IV-target genes (Figure 7B). Based on this finding, it is expected that mutating AT nucleotides in these codons to CG produces a great proportion of changes to Ala and Gly. Indeed, 24% and 11% of amino acid changes were to Ala and Gly, respectively (Figure 7C). These proportions were ∼2 times higher than those observed in *mutT*, *mutT dinB* and *mutT mutS*^β^ *dinB*. Beside the higher changes to Ala and Gly, it was evident that 55 % of mutations involved substitutions among amino acids with charged/uncharged polar and nonpolar aliphatic residues (Figure S6B). These replacements mainly included Glu to Ala, Leu to Arg, Asp to Ala and Ile to Ser. In conclusion, Pol IV preferentially mutates specific codons, favoring shifts to Ala and Gly amino acids and facilitating the change between residues with polar and nonpolar side-chains.

**Figure 7.**
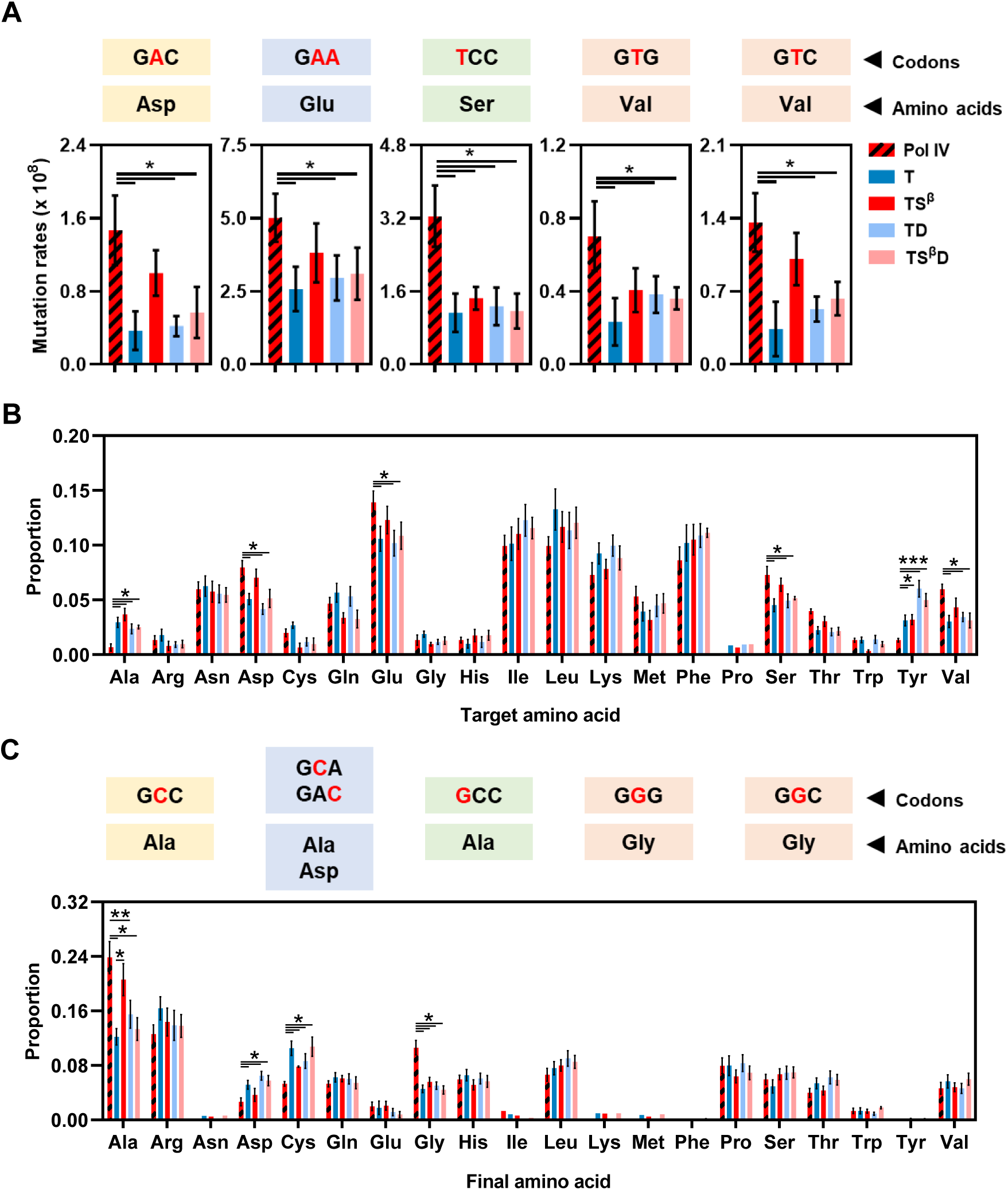
Pol IV has a bias for mutating specific codons. (A) BPS mutation rates at codons. Mutation rates per codon and generation were estimated for AT>CGs in the Pol IV-mutated genes and accumulated by the *mutT* (T), *mutT mutS*^β^ (TS^β^), *mutT dinB* (TD) and *mutT mutS*^β^ *dinB* (TS^β^D) strains. Error bars represent the upper and lower 95% confidence limits. No overlap of 95% confidence intervals indicates statistically significant differences. (B, C) Amino acid substitutions. The proportion of the target (B) and final (C) amino acids among mutations found in the Pol IV-target genes and accumulated in the T, TS^β^, TD and TS^β^D strains are plotted. Error bars represent standard deviations. The Kruskal–Wallis test was used for the statistical analysis of data.

We then investigated if Pol IV has a preference for mutating specific codon positions, which would result in either nonsynonymous or synonymous mutations. Changes in the first- and second-codon positions predominantly cause nonsynonymous changes in the coding sequence. In contrast, the third-codon position is typically degenerate, meaning that changes at this position often result in synonymous substitutions. We estimated mutation rates at each codon position for mutations promoted by Pol IV and accumulated across the four strains (Figure S7A). Mutation rates at the three codon positions were similar across all data sets. The first- and second-codon positions displayed 3- and 4-fold higher mutation rates, respectively, compared to the third-codon position. This correlates with the fact that the codon usage in *P. aeruginosa* is dominated by its high genomic GC content (67%) and the average GC content at the third position of codons is 83%^61^, making it less susceptible to AT>CG mutations. These results indicate that Pol IV does not preferentially target specific codon positions.

Furthermore, we examined whether genes targeted by Pol IV contain a higher number of Pol IV-preferred codons, which could lead to increased mutagenesis in these genes. We calculated the proportion of each codon type and the relative synonymous codon usage values for the pooled Pol IV-mutated genes (Figure S7B and Table S10). Codon usage in the Pol IV-target genes did not significantly differ from that of the entire PAO1 genome (p = 1)^61^. Likewise, the Pol IV-target genes showed a similar triplet composition to the whole-genome (p = 0.86) (Figure S7C). Thus, our analysis indicates that Pol IV-target genes do not display distinct codon usage or trinucleotide composition compared to the overall genome.

#### Pol IV-mutational genomic signatures in clinical isolates of P. aeruginosa

To investigate the potential contribution of Pol IV to bacterial adaptation, we examined single-nucleotide polymorphisms (SNPs) in the *P. aeruginosa* genome of isolates from human infections (411 genomes, including 59 from Cystic Fibrosis patients) and environmental habitats (58 genomes). Clinical and environmental isolates showed 591,087 and 467,429 SNPs, respectively, compared to the PAO1 reference genome. We first analyzed the SNP distribution across the *P. aeruginosa* genome, divided into 25 bins of 0.25 Mbp (Figure 8A and S8). SNP density per bin was calculated as the total number of SNPs in the bin divided by the number of genomes. A wave-like pattern with no significant increase in SNP density in any specific region emerged for SNPs from clinical and environmental samples. When considering different SNP types, a uniform distribution pattern was evident in transition SNPs, which constituted about two-thirds of all SNPs. Transversion SNPs were consistently distributed across the genome. However, a notably higher density was noticed specifically in the 2.50-2.75 Mbp region in the clinical isolates but not in the environmental samples. The four transversions, including AT>CG, showed 1.5- to 1.8-fold higher densities in this region relative to the overall chromosome SNP density. Particularly striking was the enrichment of transversion SNPs in the genome 2.50-2.75 Mbp segment of Cystic Fibrosis isolates, with densities reaching 2.8- to 4.5-fold higher levels. This enrichment was also evident for transition SNPs, albeit to a lesser extent (1.7- to 2.2-fold). In summary, the 2.50-2.75 Mbp region, identified in this study as a Pol IV-target region, serves as a hotspot of diversification among clinical isolates.

**Figure 8.**
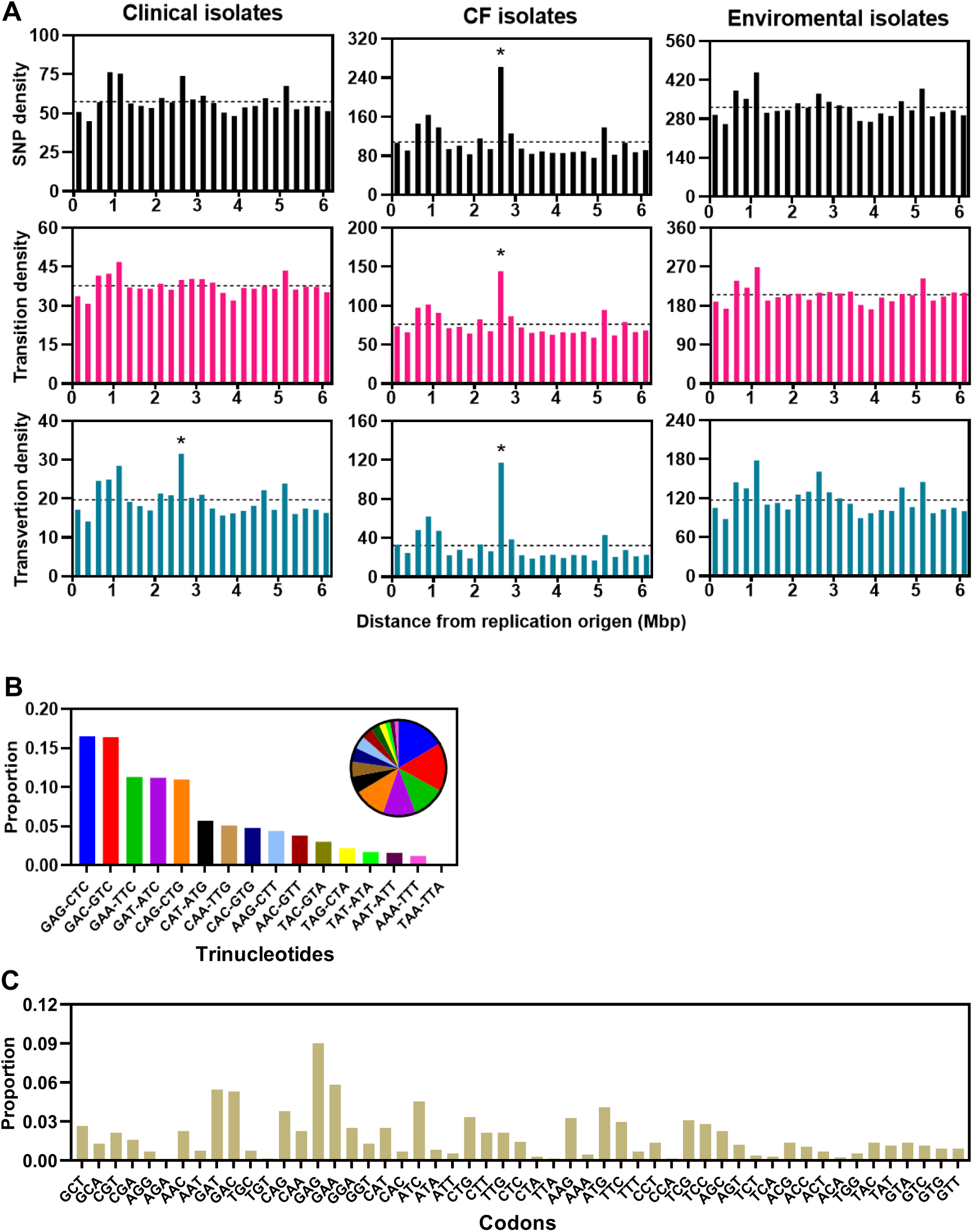
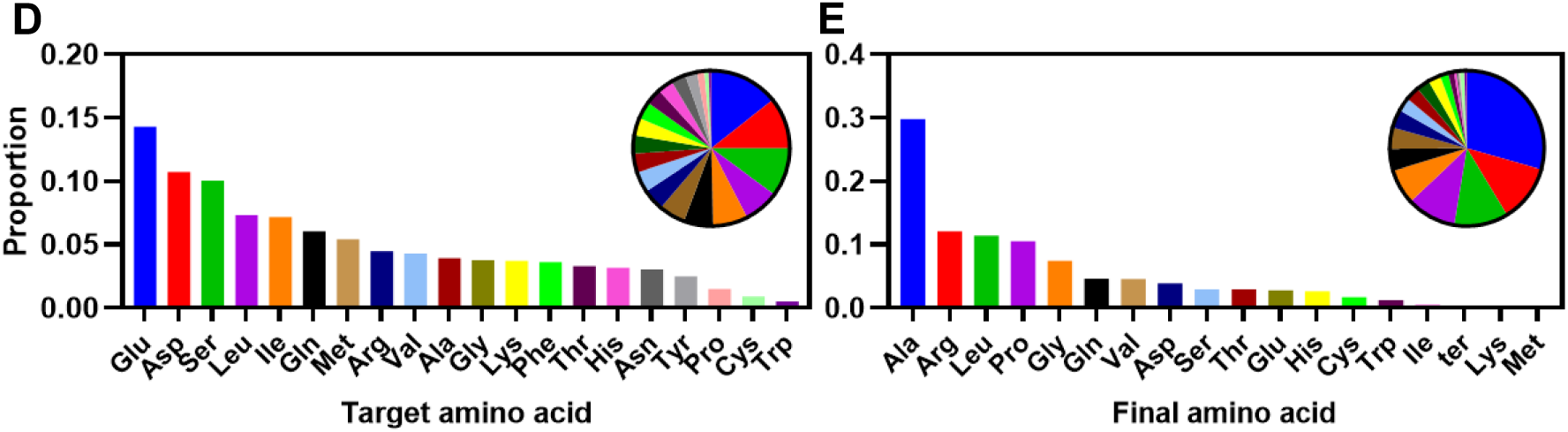
Pol IV-mutational signatures on the *P. aeruginosa* genome from clinical isolates. (A) SNP density plots for the genome of clinical, cystic fibrosis (CF) and environmental isolates. The SNP density at each 0.25-Mbp bin was calculated as the total number of SNPs in the bin divided by the number of genomes. Total SNP, transition and transversion densities were plotted as a function of distance from the origin of replication. The dotted line indicates the overall density of SNPs for the chromosome. The Iglewicz and Hoaglin’s robust test for multiple outliers (two sided test) was used to detect outlier SNP densities in the genome. Asterisks denote a significant outlier SNP density among genome regions. (B-D) AT>CG mutations in the pyoverdine pathway from clinical isolates. The proportion of mutated trinucleotides (B) and codons (C); and the proportion of the target (D) and final (E) amino acids among mutations found in the pyoverdine genes are shown.

Given that this highly variable region includes the Pol IV-target genes for the synthesis and uptake of pyoverdine, we further investigated the 1378 AT>CG SNPs identified within these genes among clinical isolates for evidence of Pol IV-mutagenesis. We analyzed the flanking sequences around AT>CGs (Figure 8B), revealing that 68% of SNPs occurred at sites with adjacent 5՚G and/or 3՚C sequences. Conversely, low occurrences (4%) were found in sequences such as T**A**A+T**T**A, A**A**A+T**T**T, A**A**T+A**T**T and T**A**T+A**T**A. In agreement with these findings, SNPs were notably enriched at codons encoding Glu (GAG and GAA), Asp (GAT and GAC) and Ser (TCG, TCC, TCA and TCT) (Figure 8C and 8D), representing 15%, 11% and 7% of AT>CG occurrences, respectively. Mutation from A to C in these codons often resulted in substitutions to Ala (Figure 8E), with 30% of SNPs leading to Ala changes.

In conclusion, our findings reveal Pol IV-mutational signatures within pyoverdine genes from clinical isolates, suggesting that Pol IV may play a role in adaptive mutation processes.

## DISCUSSION

We found that Pol IV plays a significant role in the spontaneous mutagenesis of the *P. aeruginosa* chromosome under normal growth conditions. We revealed this mutagenic effect by disrupting the non-canonical function of MutS in controlling Pol IV access to replication sites (*mutS*^β^ background)^26^. It is important to note that Pol IV-promoted mutagenesis has only been observed in this genetic context in unstressed bacteria. We previously detected mutagenesis induced by basal levels of Pol IV in the *nfxB* gene reporter^26^ and, in our current study, we also observed Pol IV action along the entire *P. aeruginosa* genome. In contrast, previous studies have not reported Pol IV-induced mutagenesis in unstressed wild type *E. coli, Bacillus subtilis and P. aeruginosa* cells using gene reporters^16,24,25^. Furthermore, Pol IV activity has been elusive across the *E. coli* genome^27^, even though the correction of replication errors by proofreading (*mutD5* strain) or MMR (*mutL* strain) was abolished^28,29^.

Mutagenesis catalyzed by Pol IV has been widely observed in bacteria under stressful conditions^16–19^. We hypothesized that this occurs because, during stress, MutS does not regulate the activity of Pol IV. Our previous research demonstrated that Pol IV-promoted mutagenesis is not constrained by the interaction between MutS and β clamp in *P. aeruginosa* cells that constitutively express the SOS response to stress-induced DNA damage^26^. This suggests that other factors may regulate MutS activity, or that MutS expression is downregulated under stress conditions. In fact, MutS levels decrease during stationary phase, carbon starvation and exposure to sublethal concentrations of antibiotics in *E. coli*^20,71^. This depletion of MutS is significant for promoting Pol IV-dependent mutagenesis induced by sublethal antibiotic levels in *E*. *coli*, *P. aeruginosa* and *Vibrio cholerae*^20^.

Our data indicate that the erroneous incorporation of the major product of guanine oxidation in the nucleotide pool underlies Pol IV-mutagenesis, resulting in AT>CG transversions. This agrees with previous biochemical and genetic evidence. Y-family Pols, including *E. coli* Pol IV, archaeal Dbh and Dpo4, and human Pol η and Pol ι efficiently incorporate oxodGTP almost exclusively opposite A *in vitro*^12,13,54,72^. It is worth mentioning that most of replicative Pols like *E. coli* Pol I and Pol II and mammalian Pol α and Pol δ can use A and C as the template base for insertion with similar efficiencies, and some of these enzymes in addition poorly incorporate oxodGTP into DNA^73–76^. The loss of Pol IV function decreases spontaneous AT>CG levels in sod/fur *E. coli* strains defective for defense against oxidative stress^13^, and impairs peroxide-induced mutagenesis in *P. aeruginosa*^11^. Incorporation of oxodGTP by Pol IV is directly involved in the lethality of bactericidal antibiotics^77^ and the increased mutation levels detected upon exposure to sub-inhibitory antibiotic concentrations^20^. Additionally, we infer from our genome-wide data that Pol IV incorporates oxodGTP and a few additional nucleotides followed by a high fidelity Pol extension of the mutagenic primer. We did not detect additional mutations in the boundaries of the AT>CG substitutions. This suggests that Pol IV does not extensively elongate the primer, leading to untargeted mutagenesis due to its low fidelity synthesis on normal template DNAs^14^. In agreement with this assumption, *E. coli* Pol IV poorly incorporates nucleotides on a primer with a terminal oxodGMP^78^.

Data from this study indicate that MutS recognizes the oxoG:A mispair and shuttle it into an appropriate accurate excision pathway to prevent AT>CG mutations. Indeed, it has been well-established that the repair factor is able to associate with oxidized DNA lesions. MutS from *Helicobacter pylori* and MutSα from *Saccharomyces cerevisiae* and human bind oxoG-containing DNA duplexes^79–81^. We propose the most likely mechanism for excision of oxodGMP involves MutS binding to β clamp, which displaces Pol IV from the clamp. This action prevents further nucleotide incorporations by Pol IV and promotes the removal of the damaged nucleotide through the exonuclease activity of a high fidelity polymerase. Supporting this hypothesis, the exonuclease activities of Pol III and Pol II have been shown to influence Pol IV-dependent mutagenesis, indicating their role in monitoring the replication products of Pol IV^82,83^. The mutator effect due to Pol IV overexpression is much higher in *E. coli* cells defective for the exonuclease ε subunit of Pol III^83^. On the other hand, inefficient extension of the primer terminus by Pol IV promotes its degradation by the proofreading activity of Pol III. Overexpression of the Pol IV Y79L mutant with a reduced ability at extending terminal ends does not result in a mutator phenotype in proofreading-proficient *E. coli* cells but induces mutagenesis in proofreading*-*deficient *dnaQ* mutant cells^84^. Another possibility is that oxoG incorporated by Pol IV triggers the mismatch repair mechanism. Certainly, a previous report showed that Pol IV errors leading to AT>CG mutations are corrected by the MMR in SOS *E. coli* mutator strains^85^. Likewise, several lines of evidence suggest a contribution of the MMR in removing oxidized lesions from DNA. *E. coli* and *S. cerevisiae* cells lacking MMR exhibit diminished mutagenesis under anaerobic growing^86,87^, while ectopic expression of the MutT homolog, MTH1, decreases mutation levels in MMR-defective mouse embryo cells and human tumors^88^. In our experiments, we cannot determine whether the mismatch repair excises oxoGs introduced by Pol IV, as MutS^β^ is proficient for mismatch correction^52^, and all strains possess a functional MMR mechanism. Additionally, we considered that the misincorporated oxoG could be removed by the DNA MutY glycosylase from the GO system. However, we ruled out this possibility because the MutY-catalyzed excision of template A in the oxoG:A mispair could lead to unwanted mutagenesis.

One of the key findings of this study is that Pol IV-dependent mutagenesis is not randomly distributed across the circular *P. aeruginosa* chromosome. We identified hotspots for Pol IV activity at specific chromosomal locations, including 0.25-0.50, 2.50-2.75, 5.00-5.25, and 5.50-5.75 Mbp. These regions have similar GC content, trinucleotide composition or codon usage as the entire genome, indicating that Pol IV targeting is not influenced by specific features in these DNA sequences. Interestingly, a notable characteristic of these chromosomal zones is their proximity to regions that pose a replication challenge. The 2.50-2.75 Mbp region, exhibiting the highest Pol IV-mutagenic activity, is located near the 3.2 Mbp region where chromosomal replication terminates^69^. The clusters of mutations at 5.00-5.25 and 5.50-5.75 Mbp are positioned between three of the four rRNA-encoding operons (*rrn*) found at 4.8, 5.3 and 6.0 Mbp, while the cluster at 0.25-0.50 Mbp is close to the fourth rRNA-encoding operon located at 0.7 Mpb^89^. Studies have shown that replication forks arrest at natural or engineered termination sites in *E. coli* and yeast^90,91^. Similarly, the highly transcribed *rrn* operons, despite being co-directional with the replication direction, provoke transcription-dependent DNA replication blockages in *B. subtilis*^92^. Furthermore, as observed in this study, chromosomal regions adjacent to these challenging-to-replicate areas are prone to increased genomic instability. Mutation rates are higher around the replication terminus region of the *E. coli*, *B. subtilis* and *P. aeruginosa* chromosomes^60,93^. Spontaneous mutagenesis of chromosomal genes is directly related to the transcription levels in *B. subtilis* and yeast^94,95^. Finally, and highlighting the connection between replication stalling and mutagenesis, the mutation rate pattern across the *E. coli* chromosome is significantly altered upon loss of the accessory replication helicase Rep, which aids replisome progression by removing DNA-bound proteins^93^.

How does Pol IV target specific genome locations for mutation? We propose that replication stalling in challenging chromosome regions, directly or indirectly, facilitates Pol IV access to replication sites, aiding replication progression. Biochemical studies show that *E. coli* Pol IV is more effective at taking over a stalled Pol III replicase due to nucleotide omission or a blocked replisome caused by a N^2^-dG adduct, helping to resume DNA synthesis^96,97^. Additionally, Pol IV molecules enrich near replisomes upon lesion stalling caused by treating *E. coli* cells with the damaging agent methyl methanesulfonate^98^. This Pol IV-recruitment at forks in the presence of a cognate lesion is also observed with forms of DNA damage that Pol IV cannot bypass, like ultraviolet light-induced lesions, as well as upon nucleotide depletion, suggesting that Pol IV localization to the replication machinery is a common response to replication stalling in *E. coli* cells^99^. Additionally, replication stalling at highly transcribed lagging-strand genes recruits the Pol IV homolog PoY1 in *B. subtilis*, facilitating replisome progression through and increasing mutation rates of these genes^94^. Although we described for the first time a preference of a TLS Pol for mutating replication challenging genome regions in bacteria, accumulating evidence suggests that these eukaryotic enzymes also support DNA synthesis through inherently difficult-to-replicate sequences. Indeed, *S. cerevisiae* REV1 and Pol ζ-dependent mutations accumulate at sites of replication fork stalling associated with non-B DNA structures^100^ while Pol ζ induces mutagenesis at telomeric repeat sequences in interstitial regions of yeast chromosomes^101^.

Another significant finding from this study is the high efficiency of Pol IV for inactivating genes. ∼56% of mutation events triggered by this TLS Pol in the *mutS*^β^ background effectively disrupted gene function under non-selective conditions. Conversely, only ∼3% of mutations induced by Pol IV-independent mechanisms in the *mutT*, *mutT dinB* and *mutT mutS*^β^ *dinB* strains resulted in gene inactivation. We obtained similar results for Pol IV-induced mutations in the *nfxB* gene selected for antibiotic resistance. We hypothesize that this feature of Pol IV-mutagenesis is attributed to the mutational signature we identified at the genome level. This DNA Pol exhibited a preference for mutating AT bases surrounded by a 5’ G and/or 3’ C. Therefore, codons containing these sequence contexts are likely hotspots for Pol IV mutations. Certainly, there was a strong bias of Pol IV towards mutating GAC, GTC, TCC, GTG, and GAA codons encoding Ser, Val, Asp, and Glu. Interestingly, these codons are commonly found in the coding regions of *P. aeruginosa* genes, comprising an average of 60% of the total codons used for each encoded amino acid. As Pol IV mutated AT nucleotides to CG in these codons, most of the changes resulted in substitutions to Ala and Gly. Both amino acids have unreactive side-chains, leading to a significant loss of protein function. Notably, Glu to Ala substitution was the most prevalent Pol IV-induced change (∼15%), suggesting that it corresponds to a Pol IV-mutation signature at the amino acid level. In conclusion, Pol IV targets specific amino acids based on its sequence context preference for mutation and changes them to the inert Ala or Gly, effectively causing protein loss-of-function alterations.

Our genomic analysis revealed that Pol IV induces mutations in specific functional categories such as virulence, motility, antibiotic resistance and chemotaxis. Genes related to the Che and Chp chemosensory systems, type IVa pili, flagellum, ABC and RND efflux pumps, membrane porins, cytotoxins and pyoverdine biosynthesis/secretion pathways were repeatedly mutated in the *mutS*^β^ genetic background. All independently evolving *mutS*^β^ lines accumulated mutations in five to six of these seven pathways. This strong enrichment of mutations across multiple lines was not evidenced for *mutT*, *mutT dinB* and *mutT mutS*^β^ *dinB* strains, which showed an average of two mutated pathways. Parallel evolution at the genomic level in the *mutS*^β^ lines under the nonselective conditions of the mutation accumulation experiment could be attributed to the fact that the genes responsible for these traits are located into the genomic regions preferred by Pol IV for mutation. The most evident examples include 70% of pyoverdine genes being situated in the 2.50-2.75 Mbp region and 83% of pili genes being located in the 5.00-5.25 and 5.50-5.75 Mbp regions. It is important to note that loss-of-function mutations in these functional pathways are common genetic adaptations found in clinical isolates of *P. aeruginosa* from chronic lung infections^35,37^. This pathogen experiences multiple convergent evolutionary changes that support its long-term survival in the hostile and stressful cystic fibrosis airways. These repeated genomic and phenotypic adaptations are often attributed to strong selection pressures in the lung environment. Therefore, it appears that Pol IV could contribute to this genetic diversification of *P. aeruginosa.* The good match between the mutation signatures found in clinical isolates and our experimentally determined Pol IV-mutation profile strongly supports an *in vivo* role for this mutagenic Pol. We detected Pol IV-mutation signatures in the highly variable 2.50-2.75 Mbp region of clinical genomes, specifically in the Pol IV-target genes for the synthesis and uptake of pyoverdine. In these hotspots of diversification, it was evident that selected AT>CG SNPs were notably enriched at sites with neighboring 5՚G and/or 3՚C and at Glu, Asp and Ser codons. Moreover, these SNPs led to a significant number of changes to Ala, with the Glu>Ala signature being prevalent among amino acid substitutions (∼12%). Remarkably, substitutions to Ala clearly inactivates protein function, as they are largely selected in the pyoverdine pathway among clinical isolates.

In sum, we conclude that Pol IV can accelerate the evolution of genes located in difficult-to-replicate regions of the chromosome, leading to the generation of highly inactivated variants as a result of its mutation features. This location-dependent mutagenesis targets commonly mutated genes in chronic *P. aeruginosa* infections, which could enhance pathogen adaptation to the harsh pulmonary environment. Further investigation is needed to ascertain whether pathoadaptive genes are located in replication challenging regions of bacterial genomes as a common mechanism for rapid evolution through the action of mutagenic DNA polymerases. Based on the results of this research, we suggest a model for how Pol IV mutagenesis could operate along genomes under stressful conditions (Figure 9).

**Figure 9.**
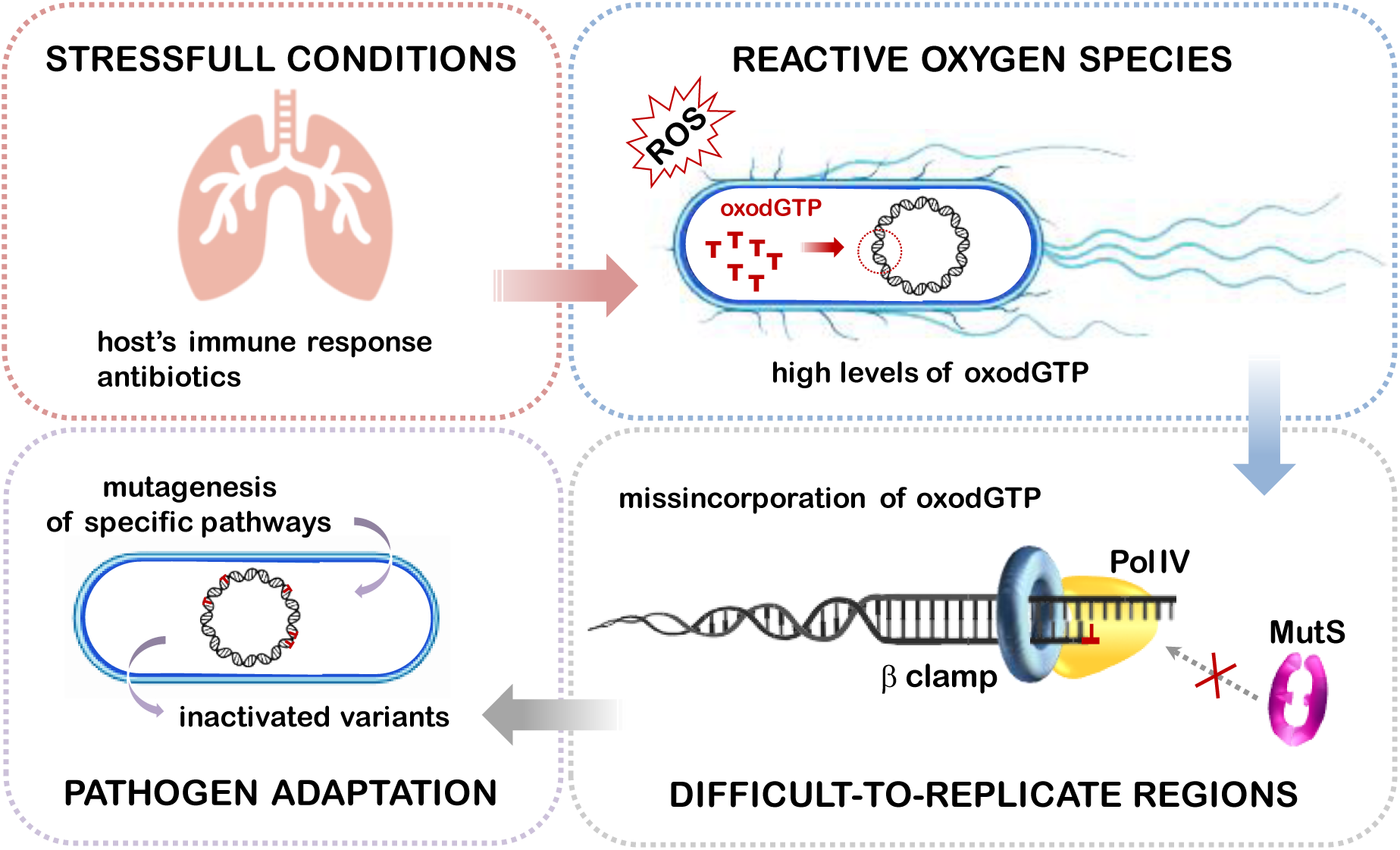
Model of Pol IV-mutagenesis across bacterial genomes. Stressful conditions, such those found in the cystic fibrosis airway environment, increase the production of reactive oxygen species (ROS), leading to the oxidation of the nucleotide pool. Fork arrest at replication termination sites and highly transcribed zones of the genome facilitates Pol IV molecules enrich near replisomes. This enzyme gains access to the processivity β clamp factor and catalyzes the erroneous insertion of oxodGTP opposite a template A surrounded by a 5’G and/or 3’C. As MutS is unable to regulate the low fidelity DNA synthesis by Pol IV under stress, these errors are fixed as AT to CG mutations. Consequently, Pol IV can drive rapid genomic evolution of genes located in difficult-to-replicate regions of the bacterial chromosome, giving rise to highly inactivated genetic variants that could enhance pathogen survival under hostile conditions.

## Notes

### Competing Interest Statement

The authors have declared no competing interest.

